# Parallel control of conjunctive and compositional representations supports dynamic task preparation

**DOI:** 10.64898/2026.01.26.701713

**Authors:** Mengqiao Chai, Iris Ikink, Stefania Mattioni, Ricardo Alejandro Benavides, Nanne Kukkonen, Mehdi Senoussi, Marcel Brass, Clay Holroyd, Senne Braem

## Abstract

People can dynamically and adaptively update task goals in the face of task uncertainty. In this fMRI study, we aimed to investigate the modulation of neural task representations that subserve such flexible task control. On each trial, we asked people to perform one of nine image categorization tasks. During task preparation, participants had to actively infer varying levels of task uncertainty and dynamically modulate their task preparation accordingly. Multivariate pattern analyses demonstrated dissociable uncertainty-driven modulations of conjunctive and compositional task representations in the right frontoparietal network. Specifically, different sub-regions showed diminished conjunctive task representations when task uncertainty increased during task preparation. At the same time, we observed an active maintenance of compositional task representations, suggesting an adaptive cognitive strategy in facilitating compositional task reconfiguration. These findings suggest a mechanism of parallel control on conjunctive and compositional representations that allows for maximizing the advantages of both representational formats, offering new insights into the neural mechanisms underpinning human flexible task preparation.

## Introduction

Navigating a multi-task environment is a control-demanding process that necessitates high cognitive flexibility (Cole et al., 2013; Garner & Dux, 2023). Such environments present two key challenges. First, real-world tasks are often compositional (e.g., assembling a piece of furniture), requiring us to dynamically integrate and decompose various task elements efficiently. Second, the rapidly evolving environment in which we live also means that tasks can be uncertain, demanding adaptive modulation of behavior based on our active inference of task uncertainty.

Classic behavioral paradigms that were typically used to study task preparation, such as task switching paradigms, are not apt to study dynamic, uncertainty-dependent changes in task preparation within a multi-task environment. To address this empirical gap, we recently developed the Task Transformation Paradigm (see Figure 1), which probes flexible task preparation under varying levels of task uncertainty. Namely, by manipulating whether the to-be-performed task could become uncertain or not within an experimental block, we can create time-dependent fluctuations in task uncertainty during task preparation. Using this paradigm, we have shown that individuals can adaptively modulate task preparation strategies in response to varying levels of task uncertainty (Chai et al., 2024, 2025). In this fMRI study, we aimed to investigate the underlying neural architecture and dynamics that subserve this flexible task preparation.

**Figure 1.**
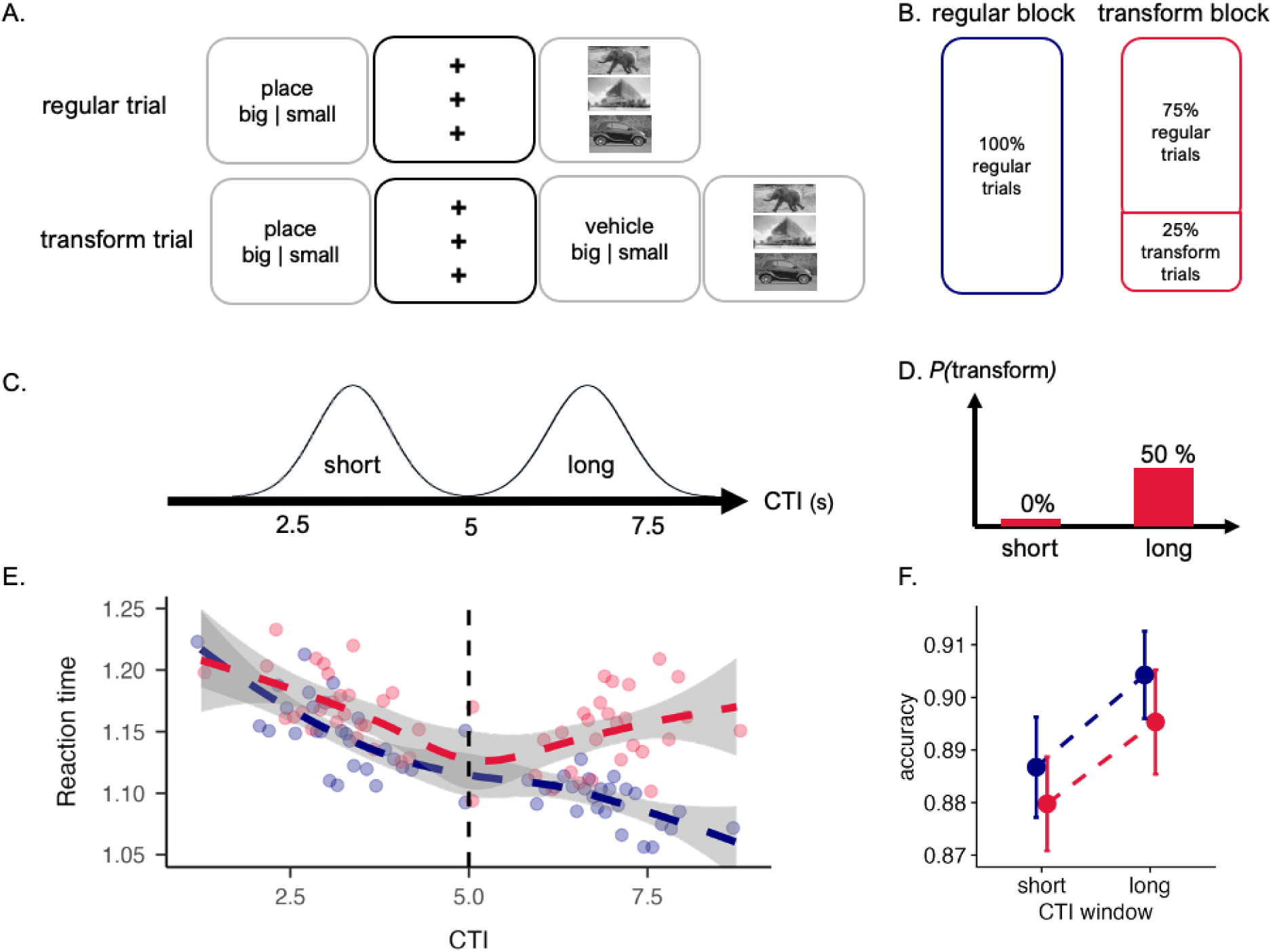
The experimental design and the main behavioral results of the Task Transformation Paradigm. **A.** A regular trial started with a task cue, and succeeded by a variable cue target interval (CTI), during which participants were asked to prepare for this task. Following this, three images representing an animal, a place, and a vehicle were displayed on screen until a participant responded by pressing the left or right key. In this example, participants were required to determine whether the image of the place could be classified as either large or small. In comparison, transform trials were featured with another task cue following the cue-target interval. In this case, participants had to perform this new task instead of the original one on the upcoming image stimuli. **B.** Two block types were included, with regular blocks including only regular trials, and transform blocks including both regular trials (75%) and transform trials (25%). Here, regular blocks can be regarded as a certain environment, and the transform blocks were the uncertain environment. **C.** The length of cue-target intervals varied across trials, by sampling from a short or long quasi-normal distribution. **D.** Critically, the length of the cue-target interval was predictive of how likely the task would transform (P(transform)). None of the trials with short cue-target intervals would transform, while half of the trials with long cue-target intervals would transform. **E.** Mean RTs of the regular trials as a function of continuous cue-target interval and block type. The colored dots represented the mean RTs across participants, which were modeled into a smooth dashed line for each block type using the loess smoothing technique. The grey band illustrates the 95% confidence interval for predictions derived from the loess regression. The purpose of loess smoothing was strictly for data visualization and not for statistical inferences. **F.** Mean accuracy of the regular trials as a function of factorial cue-target interval (short versus long) and block type. The colored dots represented the mean RTs or accuracy across participants, and the error bars denote the SE of the mean.

One of our key hypotheses regards the format of neural task representations. Prior research highlights a computational trade-off between compositional representations, which can be low-dimensional and easily generalizable, versus conjunctive representations, which are typically high-dimensional and interference resistant (e.g. Badre et al., 2021; Fusi et al., 2016; Kikumoto & Mayr, 2020; Rigotti et al., 2013). While the formation and use of conjunctive representations is likely to help in multi-task environments to prepare for different tasks while preventing interference from others, compositional task representations are more likely to benefit from recently learned or activated task components and could allow for a more flexible composition and decomposition of task sets. Therefore, given the different advantages of these two representational formats, we hypothesized that optimal task preparation could benefit from a dynamic regulation of both conjunctive and compositional task representations.

We focused our inquiry primarily within the frontoparietal network (FPN). Previous studies have typically associated the FPN with domain-general executive functions, cognitive control, and working memory (Dosenbach et al., 2006; Duncan, 2010; Fedorenko et al., 2013; Koechlin & Summerfield, 2007; Stokes et al., 2013). Critically, the FPN has been shown to adaptively represent behaviorally relevant task information in a goal-directed manner (e.g. Woolgar et al., 2011; Wisniewski et al., 2023; for a review, see Woolgar et al., 2016). In fact, due to its domain generality and flexibility in task representation, the FPN has been characterized as a flexible hub that coordinates other brain networks to subserve task performance (Cole et al., 2013; Zanto & Gazzaley, 2013), and different subregions within FPN have been observed to take on differential roles in cognitive control (e.g. Dosenbach et al., 2007; Dixon et al., 2018). Especially within the prefrontal cortex, subregions have been associated with dissociable control functions along a hierarchy, with more anterior regions being sensitive to abstract task states and more posterior regions representing more context-specific information that guides specific actions (Badre & Nee, 2018). Taken together, we expected differential neural dynamics between subregions within FPN in supporting flexible task preparation.

## Results

### Evolving task uncertainty impacted task performance

Forty-three participants’ behavior and neural activity were collected during the Task Transformation Paradigm (see Figure 1) in the MRI scanner. On each trial, participants were asked to perform one of nine image categorization tasks. Each task included two task components, which were the *stimulus type* of the target image (animal, place, or vehicle), and the *task rule* used for categorization. (i.e. big versus small, old versus young, or aquatic versus terrestrial). As depicted in Figure 1A, a regular trial started with a task cue, followed by a cue-target interval, and then image stimuli until a manual response was made. Critically, on some trials (i.e. transform trials, see Figure 1A), a task transformation would occur, meaning that the to-be-performed task would suddenly change to another task before the onset of stimuli images. This change would occur partially, meaning that only one task component would change, with the other task component remaining unchanged. If a task transformation occurs, participants have to perform this new task instead of the original one. This uncertainty in task identity required participants to regulate initial task preparation during cue-target interval when the task was being actively prepared in working memory. Another critical manipulation was that the length of cue-target interval varied across trials (see Figure 1C), with longer cue-target interval being associated with a higher likelihood of task transformations (see Figure 1D), thereby requiring participants to modulate their task preparation continuously in time.

Behavioral data indicated that participants performed well on both regular (M = 0.874, SD = 0.049) and transform trials (M = 0.93, SD = 0.054), suggesting that they did not simply disregard the need to regulate task preparation in expectation of a task transformation. Importantly, participants were tested in both a certain (i.e. regular blocks) and uncertain task environment (i.e. transform blocks), where only the latter featured transform trials (Figure 1B). To investigate differential preparatory strategies between both environments, we focused our analysis only on the regular trials in both block types. In other words, we studied how the broader context (i.e. block type), as well as its interaction with cue-target interval, affected task preparation on regular trials.

Replicating previous findings (Chai et al., 2024), a generalized linear mixed effect model on reaction time(RT) revealed a significant main effect of block type, β = -20.141, SE = 2.421, p < .001, with faster reaction times on regular trials in regular blocks (M = 1128.153, SE = 9.324) compared to transform blocks (M = 1162.265, SE = 9.963), as well as a significant interaction between block type and the length of cue-target interval, β = −5.038, SE = 1.204, p < .001, as displayed in Figure 1E. The interaction shows how participants continuously improved task performance with increasing task preparation time in a task-certain environment, as evidenced by the linearly decreasing trend of RT as cue-target interval increased in regular blocks, β = -16.543, SE = 1.608, p < .001. In comparison, this trend was weaker in transform blocks, β = -6.717, SE = 1.932, p < .001, suggesting the increased task uncertainty in transform blocks impacted task preparation. For the full statistical RT results, please see Supplementary Table S1.

Additionally, we also analyzed the accuracy of regular trials. The result from the generalized linear mixed effect model demonstrated a significant main effect of cue-target interval, β = 10.882, SE = 3.416, p = .001, with longer cue-target interval associated with higher accuracy. Unlike RT, we found no significant effect of block type, or the interaction between block type and the length of cue-target interval, ps > .149. Although non-significant, we observe a numerically higher average accuracy in regular blocks (M=0.908, SE=0.008) compared to transform blocks (M=0.885, SE=0.01). Please see Supplementary Table S2 for the full statistical results on accuracy.

### Task uncertainty activated regions within the right frontoparietal network

Turning to the fMRI results, we first aimed to identify which brain regions were activated when task uncertainty was enhanced and task transformations were expected. Therefore, we conducted a whole-brain univariate analysis by contrasting task preparation during the longer cue-target interval window between transform and regular blocks. Results showed strongly lateralized activation revealing major clusters within the FPN that demonstrated a pronounced activation in either the regular or transform blocks (for a comprehensive list of all identified clusters, see Supplementary Table S3). Specifically, right hemisphere clusters within the anterior prefrontal cortex (aPFC), lateral prefrontal cortex (lPFC), and inferior parietal lobe (iPL) were more strongly activated in transform blocks, where the to-be-performed task became more uncertain with increasing time following the task cue. In comparison, one cluster in the left lPFC responded more strongly in regular blocks, where the to-be-performed tasks were always certain.

Because participants were required to continuously regulate task preparation during cue-target interval, we further examined the specific BOLD dynamics within these significant FPN clusters using a finite impulse response (FIR) modelling approach (see Figure 2A-D). Considering the longest cue-target interval was 8.75s (see Figure 1C), and the peak delay of a BOLD response of roughly 6s (Liao et al., 2002), we estimated 10 TRs (17.8s) after task cue onset. Given that image stimuli appeared on screen immediately after the cue-target interval, we also included additional target (or “image”) onset regressors. After fitting the data to this GLM, we first selected an early perceptual region to verify whether the above described approach would mainly capture task cue-induced BOLD dynamics. As shown in Supplementary Figure S1, the BOLD dynamics of the early visual cortex clearly reflected task cue-induced activation rather than target-onset-locked activation, thereby validating our FIR modeling approach.

**Figure 2.**
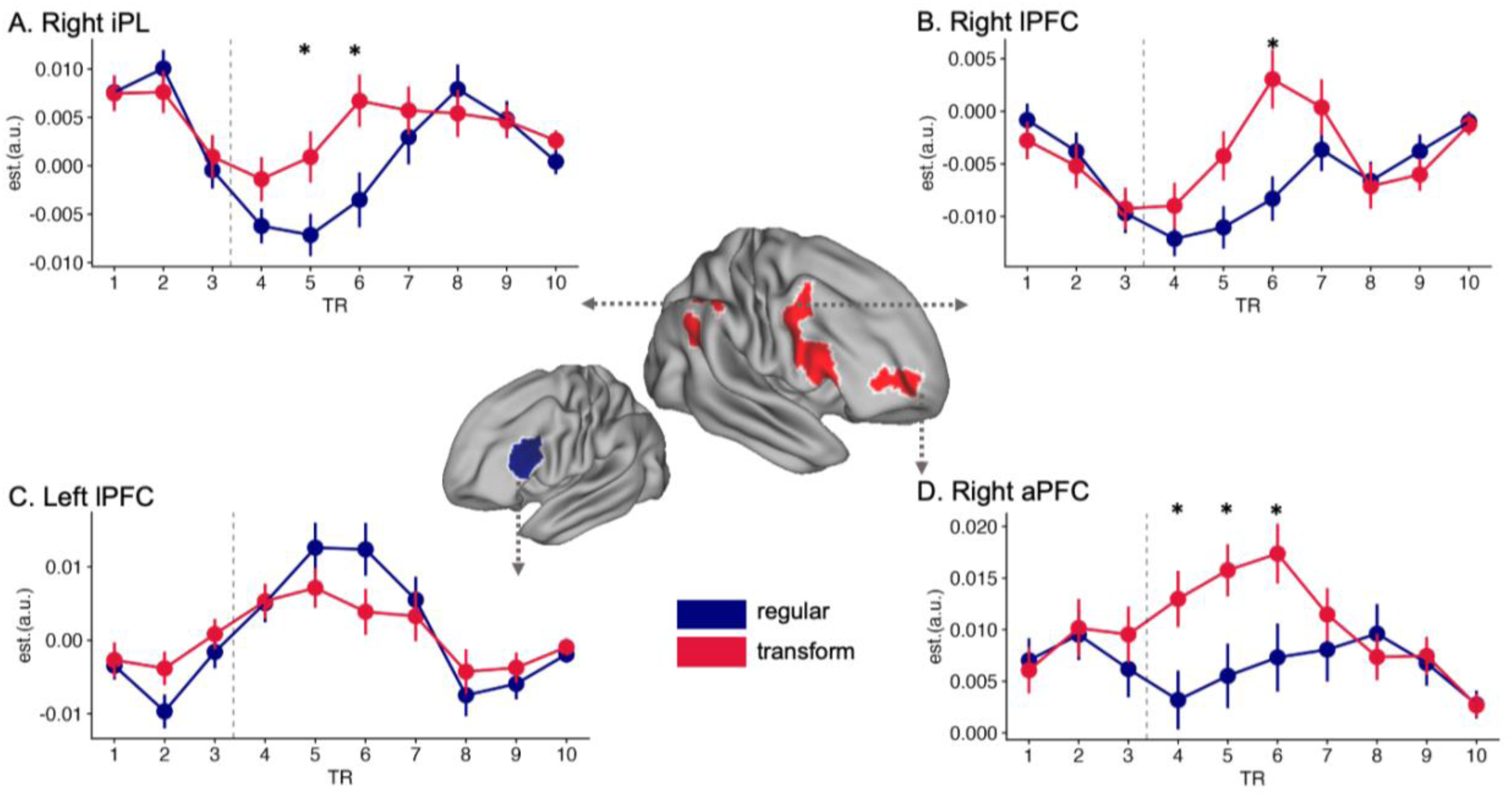
The task preparation induced BOLD dynamics across multiple brain regions, including **A.** right inferior parietal lobe (iPL); **B.** right lateral prefrontal cortex (lPFC); **C.** left lateral prefrontal cortex (lPFC); and **D.** right anterior prefrontal cortex (aPFC). These brain regions were observed in the univariate contrast that compares BOLD responses in transform blocks versus in regular blocks during long cue-target interval windows. On the surface plot, regions with crimson color were the significant clusters that showed stronger activity in transform blocks than in regular blocks. On the contrary, the region with blue color corresponds to the significant cluster that demonstrated stronger response to regular compared to transform blocks. Statistical significance was determined by firstly applying voxel level thresholding of p < .001, and then FDR cluster level correction of p < .05. The BOLD dynamics were modeled by using the finite impulse response (FIR) as basis functions in fMRI GLM analysis. Bold responses from 10 TRs (equal to 17.8 seconds) after task cue onset were estimated from this FIR GLM model. In the BOLD dynamics plots, points denote the mean model estimates across participants, and error bars denote the standard error of the mean. The dashed grey horizontal lines denote the start of long cue-target interval window (i.e. 5 seconds after cue-target interval onset). Asterisks denote TRs with significant (p <.05 without correction from t-test) differences in activation between regular and transform blocks.

Next, we examined the underlying BOLD dynamics for each significant FPN cluster identified from the whole-brain univariate analysis (see Figure 2). Consistent with our results from the univariate contrasts, the right aPFC (Figure 2D), lPFC (Figure 2B), and iPL (Figure 2A) exhibited stronger activation in transform blocks. Critically, results further revealed differential BOLD dynamics between these subregions within the right FPN. The right aPFC showed increased activation earlier during task preparation in the transform blocks compared to the right lPFC and iPL. In contrast, the right lPFC and iPL displayed heightened activity primarily after the start of the longer cue-target interval (denoted by the dashed vertical line in the plots), when a task transformation was more likely to occur. Additional exploratory uncorrected t-tests were employed to examine the difference in activation between regular and transform blocks. Results confirmed our visual inspection that right aPFC exhibited heightened activation earlier (from 4^th^ TR), as opposed to the right lPFC (6^th^ TR) or right iPL (5^th^ TR). Although we should be cautious in making strong conclusions about the temporal sequence of information processing due to low temporal resolution of the BOLD signal, both visual inspection and these exploratory statistical tests suggest the right aPFC was activated earlier, following task cue encoding, and this activation sustained longer. In comparison, the more posterior right lPFC and iPL showed a slightly more delayed ramp-up in activity in response to increased task uncertainty within the cue-target interval.

To assess the robustness of our findings on the whole-brain univariate contrast, we also adopted other GLM approaches (see alternative GLMs in Methods). All identified brain clusters were highly overlapping (see Supplementary Figure S2 & S3), especially within the FPN, confirming the robustness of our univariate results, and highlighting the key involvement of the right FPN in responding to evolving task uncertainty in a multi-task environment.

### Task decoding of conjunctive task representations revealed uncertainty-driven modulation on representational strength

To further investigate the role of right FPN in subserving flexible task preparation, we examined the dynamics in task representational strength through decoding analyses. Specifically, we decoded conjunctive task information in each ROI within the right FPN as a function of block type and cue-target interval (see Methods for more detailed decoding set-up). The conjunctive task representation is defined as the task identity integrating both stimulus type and task rule dimensions. In other words, these decoding analyses quantified how much the voxel patterns of each of the nine tasks could be separated in voxel space. The easier these patterns can be separated, the higher the decoding accuracy should be, indicating stronger task representations.

The ROIs used for decoding were defined independently (see ROI definitions from the Methods section of multivariate analysis). We first set out to test the overall decodability in FPN given the difficulty of task decoding in fMRI data, especially in the prefrontal cortex (Bhandari et al., 2018). As a result, we found significantly above-chance decoding accuracy in FPN (see Figure 3A), β = 0.0085, SE = 0.0016, p < .001, confirming the overall task decodability in FPN. Critically, when comparing to a task-neutral network (i.e. the early auditory network), we found significantly higher decoding accuracy in the FPN, *t*(42) = 4.2598, *p* < .001, and no above-chance decoding performance in this task-neutral network, β = - 0.0014, SE = 0.0017, p = .413, further illustrating the specificity of the FPN in representing task information.

**Figure 3.**
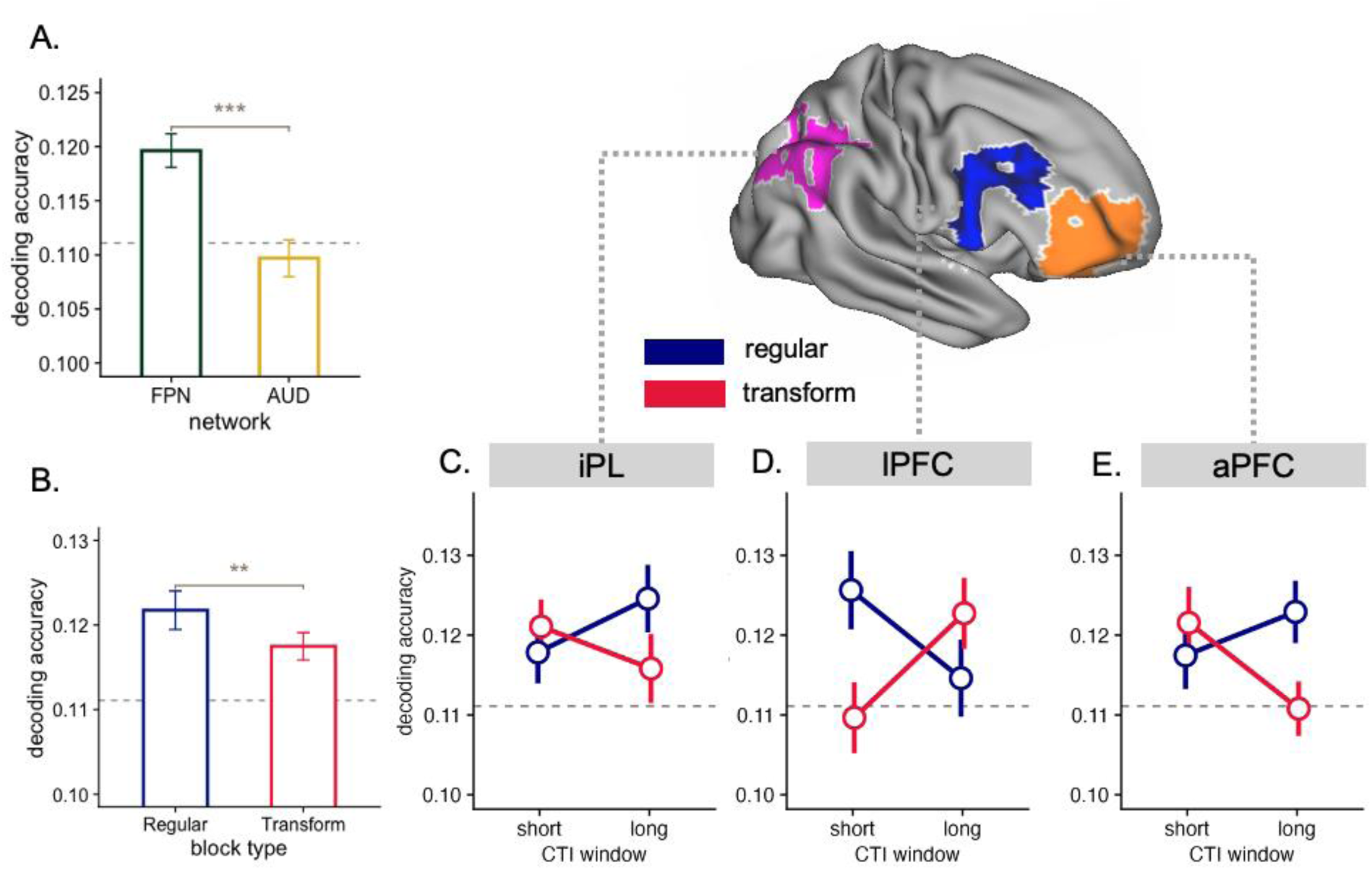
The results from conjunctive task decoding of both regular and transform blocks within FPN. **A.** The mean decoding accuracy in the frontoparietal network (FPN) and the early auditory cortex (AUD). Both networks were independently defined by using MMP atlas (Glasser et al., 2016). The FPN was additionally selected from the MD system identified by Assem et al. (2020). Here, the significant above-chance decoding in FPN is contrasted with the below-chance decoding in AUD, confirming the specificity in representing task information within FPN. **B.** The decoding accuracy in regular blocks was significantly higher than transform blocks, indicating that participants represented task information stronger in a task-certain environment than in a task-uncertain environment. **C-E.** The decoding accuracy as a function of interaction between cue-target interval window and block type for each ROI within right FPN, including **C.** iPL, **D.** lPFC, and **E.** aPFC. Both aPFC and iPL demonstrated the interaction that the representational strength (indexed by the decoding accuracy) of conjunctive task increased during cue-target interval when participants were certain about the to-be-performed task in regular blocks, meanwhile the representational strength decreased during cue-target interval in transform blocks since long cue-target interval window was predictive of a potential task transformation. In comparison, the lPFC demonstrated a somewhat opposite interaction by showing more pronounced differences in representational strength between block types during short instead of long cue-target interval window. Bar and dots in the plots denote the mean decoding accuracy across participants, and error bars denote the SE of the mean, grey dashed lines denote the chance-level for conjunctive task decoding (i.e. 0.1111). * denotes p < .05, ** denotes p < .01, *** denotes p < .001.

We expected that the inclusion of transform trials in transform blocks would generally weaken conjunctive task representations compared to regular blocks, leading to lower task decoding accuracy in transform blocks compared to regular blocks. Indeed, this is what we found (see Figure 3B), β = -0.0020, SE = 0.0007, p = .006. We further expected to see dynamic changes in task representations similar to our main behavioral findings (see Figure 1E). That is, in regular blocks we expected the strength of task representation to increase during the cue-target interval, as suggested by continuous improvements in task preparation when the to-be-performed task was certain. In contrast, we expected decoding accuracy to either decrease or increase less in transform blocks, as participants may need to adaptively down-regulate a conjunctive task representation when a task transformation is imminent.

As displayed in Figure 3, both the right aPFC (Figure 3E), and iPL (Figure 3C) showed or hinted at this expected pattern in task decoding accuracy, as suggested by the significant interaction between cue-target interval window and block type in the right aPFC, β = 0.0041, SE = 0.0017, p = . 0199, and non-significant but numerically similar in the right iPL, β = 0.0030, SE = 0.0016, p = .0536. Interestingly, the lPFC demonstrated a different interaction, β = -0.006, SE = 0.0021, p = .0039, showing a stronger difference in decoding accuracy between regular and transform blocks in the shorter cue-target intervals rather than longer cue-target intervals, suggesting a differential role played by right lPFC in uncertainty-driven task preparation. An overall analysis comparing the cue-target interval window and block type interaction between right aPFC, iPL, and lPFC (aPFC + iPL versus lPFC) indeed yielded a significant difference, β = -0.0382, SE = 0.0096, p < .001, supporting the regional differences in uncertainty-dependent modulations of conjunctive task representations over time. Please see Supplementary Figure S4 for results for all ROIs within bilateral FPN, and Supplementary Table S4 for the full statistical results. We also conducted the above analysis by using an alternative ROI definition scheme (see ROI definitions from the Methods section of multivariate analysis for more details) and yielded highly comparable results (Supplementary Figure S5), thereby confirming the robustness of the above-presented results on conjunctive task decoding.

### Compositional task representations were actively maintained in right aPFC and iPL despite diminished conjunctive representation

Next, we quantified the strength of compositional task elements that jointly define the task to study their potential complementary role in supporting flexible task preparation (see compositional task decoding in our Methods section). For instance, a task can be defined as “animal” in the stimulus type dimension, and “small | big” in the task rule dimension. Together, they formed one conjunctive task out of nine. Now, rather than decoding this conjunctive task representation, we can also decode both compositional task elements separately. The decoding accuracy of these compositional elements was indicative of how strongly the task was encoded in a compositional format. Previous studies already shed light upon differential dynamics of conjunctive and compositional representations, and how they may support people’s task performance (e.g. Kikumoto & Mayr, 2020). Here, we aimed to examine whether people can leverage the distinct advantages of both representational formats to facilitate flexible changes in task preparation when preparing for task uncertainty (e.g. Fusi et al., 2016; Badre et al., 2021).

We first examined the overall decodability of compositional task elements in the FPN. As a result, we observed overall above-chance decoding accuracy on both task dimensions (see Figure 4A), including stimulus type, β =0.0268, SE = 0.0056, p < .001, and task rule, β = 0.02, SE = 0.0053, p = .001. These results indicated that, in addition to the conjunctive format, the task information was also represented in a compositional format within FPN. Moreover, we observed a higher decoding accuracy on the dimension of stimulus type compared to task rule, β =-0.0034, SE = 0.0016, p = .031, indicating that, during task preparation, the FPN showed a slightly stronger encoding of stimulus type dimension compared to task rule dimension.

**Figure 4.**
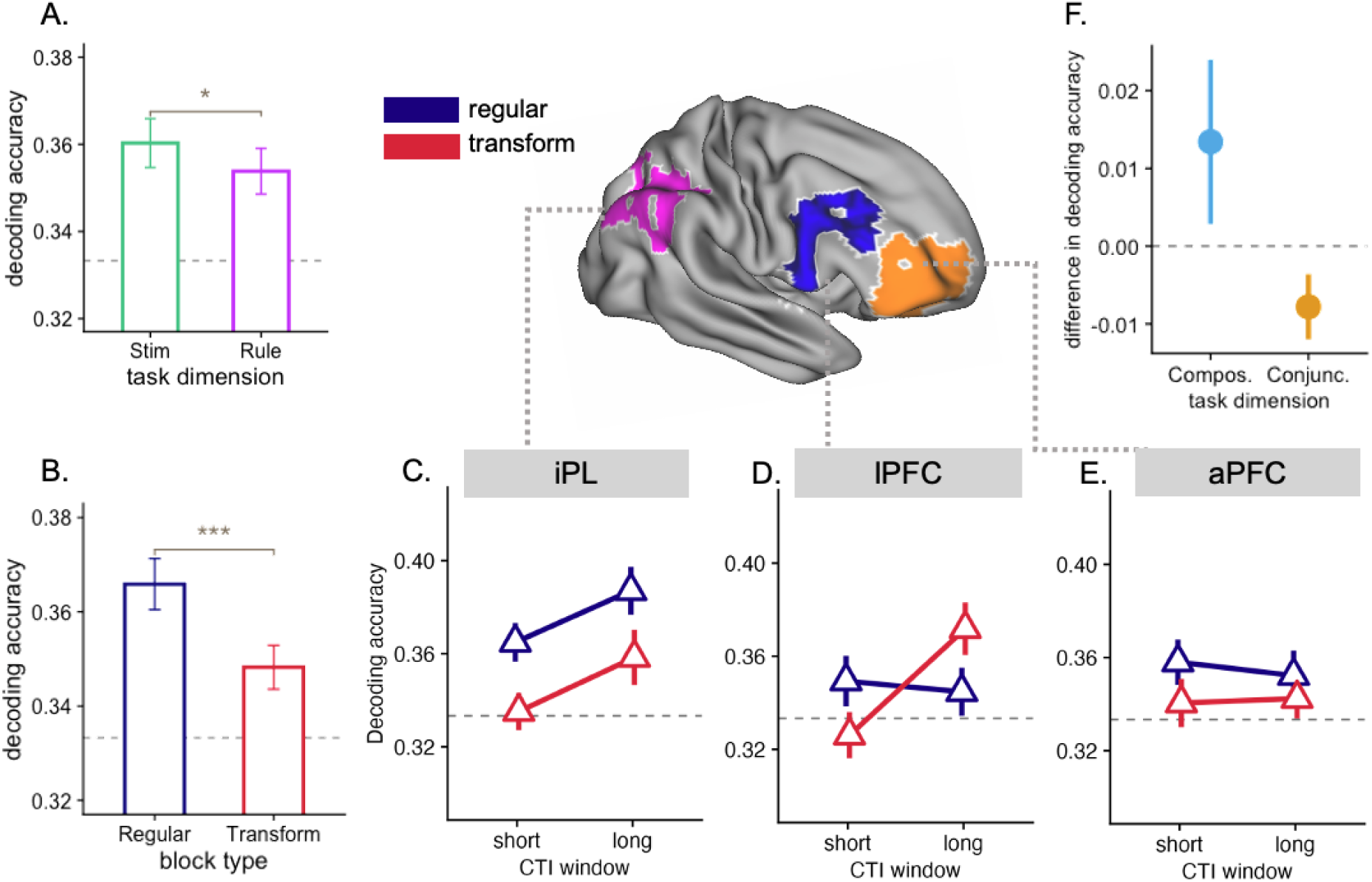
The results from compositional task decoding of both regular and transform blocks within FPN. **A.** The overall above-chance level decoding across two task dimensions, including stimulus type (Stim) and task rule (Rule), with higher decoding accuracy of stimulus type dimension than task rule dimension **B.** The compositional decoding accuracy in regular blocks was significantly higher than transform blocks, indicating stronger compositional representation in a task certain environment than in a task uncertain environment. **C-E.** The compositional decoding accuracy as a function of interaction between cue-target interval window and block type for each ROI within right FPN, including **C.** iPL, **D.** lPFC, and **E.** aPFC. Compositional representation of these regions unanimously demonstrated sustained or increased activation during cue-target interval in transform blocks. **F.** The comparison of cue-target interval effect (long cue-target interval window – short cue-target interval window) in transform blocks between compositional (Compos.) and conjunctive (Conjunc.) decoding accuracy in right iPL and aPFC. Result confirmed that these two ROIs exhibited overall increasing compositional decoding accuracy during cue-target interval (positive on the y axis), in comparison to decreasing conjunctive decoding accuracy during cue-target interval (negative on the y axis). Bars and dots in the plots denote the mean decoding accuracy across participants, and error bars denotes the SE of the mean, grey dashed lines of **A-E** denote the chance-level for compositional task decoding (i.e. 0.3333).

Next, we further inspected whether the overall task uncertainty impacted the strength of compositional task representations. Similar to conjunctive task decoding, we observed higher compositional decoding accuracy in regular blocks compared to transform blocks (see Figure 4B), β = 0.0082, SE = 0.0016, p < .001.

When testing the interactions between cue-target interval window and block type in the ROIs within right FPN, only the lPFC yielded a significant result (see Figure 4D), β = -0.0126, SE = 0.0043, p = .0036. Interestingly, this interaction was similar to that observed in conjunctive task representations, suggesting converging representational dynamics for conjunctive and compositional task representations in the right lPFC.

In contrast, no interaction was observed in either the right aPFC or right iPL (see Figure 4C, E), ps > .59. As discussed above, both of these regions displayed decreasing representational strength in conjunctive task representations during cue-target interval of transform blocks (see Figure 3C, E) to facilitate the potential upcoming task transformation. However, we did not observe a decreasing compositional representational strength in either of these two regions, implying that compositional task representations sustained throughout cue-target interval despite increasing task uncertainty in the longer cue-target interval window. A post-hoc t-test that compared the effects of cue-target interval effect (the decoding accuracy between cue-target interval long window and cue-target interval short window) between compositional and conjunctive decoding in these two regions indeed demonstrated a significant difference (see Figure 4F), *t*(42) = 2.2627, *p* = .014, supporting our observation of a concurrent decreased conjunctive representation but sustained compositional representation in transform blocks within the right aPFC and iPL. Please see Supplementary Figure S6 for the results of compositional decoding for each task dimension separately, and Supplementary Table S5 for the full statistical results.

### Task uncertainty led to noisy representation

Finally, to test whether task uncertainty impacted overall representational consistency in these different regions, we also computed the distance (i.e. Spearman’s correlation) between voxel patterns of each (conjunctive) task representation based on the premise that the distance between voxel patterns were indicative of how consistent the representation was across runs. That is, a smaller distance corresponds to higher representational consistency.

We reasoned that higher task uncertainty could also disrupt the overall consistency of neural pattern responses in the FPN, potentially reflecting a strategic lowering of neural gain to allow a more flexible switching between task representations (e.g., Musslick & Cohen, 2021), leading to lower representational consistency. Indeed, we observed a significantly lower representational consistency in transform blocks than regular blocks (see Figure 5A), β = 0.0077, SE = 0.0011, p < .001. In addition, we observed higher representational consistency in long compared to short cue-target interval windows (see Figure 5B), β = -0.0231, SE = 0.0011, p < .001, and no significant interactions between block type and cue-target interval (see Figure 5D, E, F), ps > .317.

**Figure 5.**
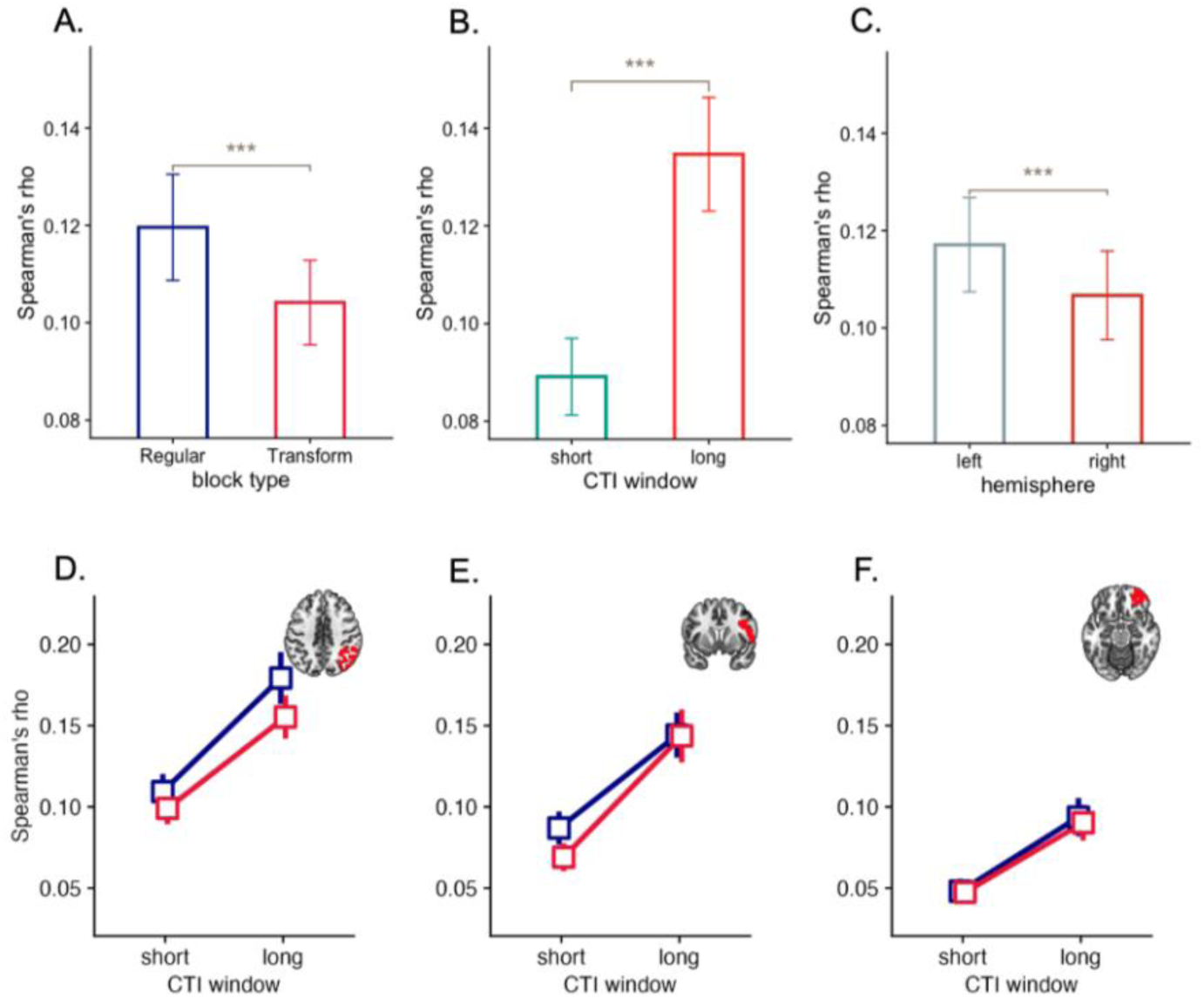
The results of representational consistency analysis. **A.** the mean representational consistency, which was measured by Spearman’s correlation between voxel pattens of a certain task across runs, in regular and transform blocks. A higher representational consistency was observed in regular than in transform blocks. **B.** The mean representational consistency in short and long cue-target interval windows. A higher consistency was found in long than in short cue-target interval window. **C.** The mean representational consistency between left and right FPN. There was a higher consistency in the left compared to the right FPN. The mean representational consistency for each ROI within right FPN as a function of interaction between cue-target interval window and block type was illustrated in **D.** iPL, **E.** lPFC, and **F.** aPFC. No clear interaction effect was observed across these ROIs. Bar and dots in the plots denote the mean representational consistency, measured by Spearman’s rho, across participants, and error bars denote the SE of the mean.

We did find significantly lower representational consistency in the right hemisphere compared to the left hemisphere (see Figure 5C), β = 0.005, SE = 0.0011, p < .001, which was in line with our results from univariate contrast showing that right FPN responded strongly to task uncertainty. For the full (statistical) results of representational consistency for all ROIs within bilateral FPN, please see Supplementary Figure S7 and Table S6.

## Discussion

We investigated the underlying dynamics of neural task representations within the frontal-parietal network (FPN) that support uncertainty-driven flexible task preparation. We used a newly developed paradigm where participants dynamically changed task preparation based on their active inference of task uncertainty (Chai et al., 2024, 2025). Using fMRI, we demonstrated how different regions in the right FPN subserve these dynamic changes in task preparation through a flexible reweighting of conjunctive versus compositional task representations. The right iPL and aPFC in particular, selectively downweighed conjunctive task representations in the face of increasing task uncertainty, while retaining information about the compositional task elements to allow for flexible task reconfiguration.

The active recruitment of the FPN during task preparation fits with previous literature that has characterized the FPN as a domain general network typically recruited when performing cognitively demanding tasks (Fedorenko et al., 2013; Braver et al., 2021; Assem et al., 2020). Notably, activity in the FPN was strongly lateralized, with greater activity in the right FPN under conditions of high task uncertainty (e.g., transform blocks). The right hemisphere of the human brain has been described as being “*specialized for detecting and responding to unexpected stimuli in the environment*.” (MacNeilage et al., 2009). Previous literature has also associated right frontal-parietal regions, mainly including iPL, lPFC, and superior temporal gyrus (STG), with detecting behaviorally relevant or novel stimuli (Corbetta &Schulman, 2002), arousal (Sturm et al., 1999), and orientating attention, especially guided by cognitive control (Shulman et al., 2009). Given their frequently observed co-activation, these regions have been theorized to form the ventral attentional network that encompasses the above-mentioned cognitive functions.

In addition to right iPL and lPFC, we also found stronger activation in the right aPFC. The aPFC or, more precisely, rostral lateral PFC (BA 10) is a highly debated brain region in terms of its functional specificity. This region has been characterized to sit on top of a rostro-caudal hierarchy within the frontal lobe, with its key function being the representation of abstract schematic control (Badre & D’Esposito. 2009; Badre & Nee, 2018). Additionally, rostral lateral PFC has been regarded as crucial in exploring alternative or competing goals (e.g. Mansouri et al., 2015). It has also been argued that this region is critical to maintaining alternative task sets in a multi-task environment (Benoit., 2008). Mansouri and colleagues (2017) posited that rostral lateral PFC serves as the key region to support directed exploration by monitoring alternative goals and redistributing cognitive resources between them (also see Koechlin et al., 1999; Dreher et al., 2008). Consistently, participants in our paradigm also face this challenge because they have to make a choice to either keep preparing for the current task (*exploitation*) or prepare to switch to an unknown alternative task (*exploration*). Taken together, the activation of the right aPFC may broadly reflect this weighing of these two competing goals during task preparation, especially when the cue-target interval was getting longer in transform blocks.

The rostral PFC has also been proposed to be a key region in re-orienting one’s control state from stimulus-oriented to stimulus-independent based processing according to the gateway hypothesis (Burgess et al., 2007; Gilbert et al., 2006), highlighting the critical role of rostral PFC in transitioning between external and internal attention. In our study, participants were asked to use the cue-target interval to actively prepare for the task based on the task cue. This cognitive process was initially stimulus (i.e. task cue) oriented. However, when the cue-target interval was getting longer in transform blocks, participants had to infer a change in task uncertainty, and adjust their preparation strategy accordingly. This process is self-generated without external cueing (stimulus-independent). Thus, this shift in cognitive control strategy could also contribute to the reason why we observed a boost of activation in rostral PFC.

Interestingly, a closer investigation of the different time-varying activation dynamics indicated an earlier responsiveness of the aPFC compared to the lPFC and iPL, both of which exhibited more pronounced activity towards the longer cue-target interval. These activation dynamics of lPFC and iPL nicely mapped onto the evolution of task uncertainty during task preparation. Namely, activation was enhanced when task uncertainty increased during the cue-target interval. In comparison, right aPFC enhanced its activation more early on during task preparation. This result is in line with previous studies showing that aPFC was involved when the temporal context needed to be closely monitored to reach one’s goal (e.g. Nee et al., 2014; Volle et al., 2011). Here, too, the temporal information (i.e. time elapsed during cue-target interval) was more critical and informative to monitor task uncertainty in transform blocks compared to regular blocks. Therefore, this early activation in right aPFC may have reflected a stronger monitoring of temporal context.

Moving beyond its broader role in monitoring task uncertainty, an important aim of this study to get a deeper understanding of the role of the right FPN in supporting flexible task preparation by quantifying the strength of neural task representations. To investigate whether conjunctive and compositional task representations were subject to dissociable modulations during task preparation under varying levels of task uncertainty, we conducted both conjunctive and compositional task decoding. Both right aPFC and iPL demonstrated an adaptive modulation of conjunctive task representation, with increasing representational strength during task preparation in a task-certain environment, and decreasing representational strength when expecting a potential task switch in a task-uncertain environment. These results align well with previous studies that highlighted the highly flexible nature of FPN in representing behaviorally relevant information (for a review, see Zheng et al., 2024), which is also supported by findings showing adaptive information encoding depending on task demand (Woolgar et al., 2011; Wisniewski et al., 2023), reward contingency (Hall-McMaster et al., 2019; Etzel et al., 2016), and task history (Qiao et al., 2017). Our results extended this line of research by showing that a modulation of such representational strength can occur within a short interval, which resonates with the notion of a dynamic coding scheme of working memory (WM) in PFC. This theoretical framework on WM emphasized the rich time-varying dynamics during WM maintenance and how it can offer computational benefit in supporting optimal behaviors (Stokes et al., 2013; Stroud, Duncan & Lengyel, 2024). However, the BOLD signal delay and the low sampling frequency (TR=1.78s) render fMRI not the best tool for studying fast neural dynamics. Thus, future studies could employ EEG or MEG recording to further unveil this dynamic process.

Importantly, in contrast to the decreasing representational strength in this conjunctive format, the representational strength of the compositional elements was maintained across time, implying that these two representational formats were regulated in parallel. In our paradigm, task transformations occurred only partially, meaning that only one of two task components was transformed. Therefore, although an expected task transformation implied that a conjunctive task representation could become obsolete, one of two compositional elements would still be relevant as part of the newly transformed task. In this context, the optimal strategy involved simultaneously downregulating conjunctive task representations while actively maintaining compositional ones, which is exactly what we observed in the right aPFC and iPL. These results are in line with previous works on representational dimensionality that demonstrated how neural task representations embed both high- and low-dimensional information, allowing for both an efficient read-out for downstream processes and abstractions across tasks in order to generalize (Bernardi et al., 2020; Xie et al., 2022). Our findings extend this line of research by suggesting that high (conjunctive) and low-dimensional (compositional) task representations can be controlled independently in an adaptive and goal-directed manner. By representing tasks in both conjunctive and compositional formats, the advantages of both representational formats can be leveraged, going beyond the notion that a trade-off is required between high and low dimensional task representations (Badre et al., 2021).

In comparison to the right aPFC and iPL, the right lPFC exhibited opposite dynamics in conjunctive task representation. Namely, the lPFC showed stronger conjunctive task representations in certain as opposed to uncertain environments from very early on during task preparation, even though differences in task (un)certainty between these two environments arose only during long cue-target interval window. Although speculative at this point, as we did not predict this result, it is possible that in the task-certain environment (i.e. regular blocks), individuals know exactly which task they will need to perform during task preparation, thereby allowing for a more efficient and focused initial encoding of task-related information immediately after task cue onset. Therefore, the lPFC, which is often thought to prepare a more procedural task format closer to the concrete action plans, can immediately engage in forming a task-ready conjunctive task representation. In comparison, in the task-uncertain environment (i.e. transform blocks), participants might initially be reluctant to form task representation due to higher task ambiguity. However, as time progresses, they might gradually form conjunctive task representations in preparation for the subsequent stimulus onset. This progressive preparation could lead to increased lPFC task encoding, but only later in time during the long cue-target interval window. Notably, the right lPFC also demonstrated a similar interaction pattern in compositional task representation, suggesting a consistent dynamic modulation on task representation across representational formats in this region.

Finally, there has been a recent increasing interest in studying the variability of neural signals and its function in human cognition. It has been proposed that neural variability could play a key role in cognitive flexibility (Uddin, 2021; Dajani & Uddin, 2015), especially regarding uncertainty-driven behavior (Waschke et al., 2021; Kosciessa et al., 2021). Specifically, this variability could be driven by stochastic resonance, or stochastic facilitation, which has been observed to increase the responsiveness in the neural system in both theoretical and experimental neuroscience (Garrett et al., 2011; Ward & Greenwood, 2016; McDonnell & Ward, 2011). In our results, we observed that voxel pattern consistency was decreased in uncertain task environments (i.e. transform blocks). One interpretation is that this decreased consistency was driven by less strong task representations, as supported by our results of both conjunctive and compositional task decoding. Another non-mutually exclusive possibility is that neural noise or strategic variability was functionally injected into the FPN during task preparation when a transformation was expected in order to facilitate the subsequent encoding of new task cues via the mechanism of stochastic facilitation. However, previous works on neural variability typically recorded electrophysiological signals given its high temporal resolution and sampling frequency. Therefore, we believe a more vigorous test of such a hypothesis would require techniques like EEG or MEG to further unravel the potential functional role of neural variability in supporting uncertainty-driven behavior.

In conclusion, our results provide insight into how the frontoparietal network supports flexible task preparation in the face of increasing task uncertainty. The right FPN showed parallel control of conjunctive and compositional task representations, consistent with an adaptive strategy that maximizes the advantages of both representational formats.

## Methods

### Participants

47 young adults (age between 18 and 35) were recruited for this study. All participants were right-handed, had good understanding of English, and good eyesight (including corrected normal vision), had no neurological history or took any anti-depressant. Participation was compensated by payment of €60 in total for two scanning sessions and a short online practice.

Prior to the data collection, ethical approval was acquired from the Committee of Medical Ethics of University Hospital of Ghent, Belgium (UZ Gent). Informed consent were acquired from each participant at the beginning of each scanning session. Participants also had to fill out a pre-scanning and post-scanning checklist to ensure MR safety for each scanning session.

Data from two participants were excluded due to drop-out after the first scanning session. Data from another two participants were further excluded due to their near chance-level performance on the task, as well as their post-scanning self-reporting on their heavy drowsiness during the scanning. After the above exclusion, data from 43 participants (24 females, 19 males, mean age = 27) remained for analysis.

### The task procedure

Participants were asked to complete a variation of Task Transformation Paradigm (Chai et al., 2024), which was designed to investigate dynamic changes in task preparation. In this image categorization task, participants had to categorize images based on both the stimulus type the image belonged to (either animal, building, or vehicle), and a task rule that dictated the property (either size, age, or habitat) on which the image should be categorized. By permuting these two compositional task dimensions, nine different tasks were created. And for each trial, participant had to first encode the task cue, then actively prepare for this task during cue-target interval, and then categorize the relevant image correctly by pressing either the left or right key based on the task cue (see Figure 1A).

On transform trials, a new task cue could be displayed following the cue-target interval. The onset of the new task cue required participants to categorize the incoming images based on this new task cue instead of the original one that they were preparing for during the cue-target interval (see Figure 1A). Thus, the to-be-performed task could be uncertain during task preparation. Moreover, the length of cue-target interval varied across trials between 1.25s to 8.75s (M = 5.04s). Cue-target intervals with a short length followed a quasi-normal distribution with M = 3.16s, and SD = 0.76s. The long cue-target intervals also followed a quasi-normal distribution with M = 6.92, and SD = 0.76s (see Figure 1A). Importantly, the length of the cue-target interval was indicative of the likelihood of task transformation: only long cue-target intervals were associated with a 50% likelihood of transformation (see Figure 1D). Therefore, it required participants to keep track of time elapsed during task preparation and to adjust their task preparation dynamically based on this phasic task uncertainty within the cue-target interval.

Importantly, we created two environments with different levels of task uncertainty. In one environment (i.e. regular blocks), no task transformation could be expected. Thus, participants were always certain about the task they were going to perform during the cue-target interval. In comparison, in another environment (i.e. transform blocks), half of trials with a long cue-target interval involved task transformation, requiring participants to manage task uncertainty throughout the whole block (see Figure 1B).

Before the scanning session, each participant went through the instructions online, which was programmed in JsPsych version 6.1 (de Leeuw, J. R., 2015). During this, participants were also required to categorize each image stimulus correctly, and went through 20 trials of regular condition and 6 trials of transform condition to make sure that they could be able to perform the task relatively well from the very start of the scanning session.

The experiment itself was created and presented in PsychoPy 2024.2.1 (Peirce et al., 2019). Each participant went through two scanning sessions at the University Hospital of Ghent (UZ Gent) separated by one to three days. Each scanning session contained 4 runs of same block type, meaning that participants would finish 4 runs of one block type in the first scanning session, and finish another 4 runs of the other block type in the second scanning session. The starting block type was counterbalanced across participants (22 of them started with regular blocks, and 21 of them started with transform blocks) to avoid the potential confound of time-on-task effect on a group level. Each block consisted of 54 trials, with each of 9 tasks being repeated 6 times in a random order, with the only constraint that there could be no task repetition between two consecutive trials. Half of the trials within a block were trials with short cue-target intervals, and the other half were with long cue-target intervals, with short and long trials being randomly interleaved. Participants were allowed to take short breaks between blocks. Each scanning session lasted around 75 to 90 minutes in total.

### Behavioral data analysis

Behavioral data analysis mainly focused on the reaction time (RT). Data processing and analysis were performed using R (R Core Team, 2017). Consistent with the data exclusion procedures of previous experiments (e.g. Chai et al., 2024), participants with less than 60% accuracy in regular or transform trials were excluded, as were those whose average RT or error rate exceeded ±2.5 SD from the group mean. Error trials, and trials following errors, faster than 200 ms, or with RTs ±2.5 SD from the individual mean were also removed from the analysis. After data exclusion, the RT data were fit with a Gamma distribution by using generalized linear mixed effect models (glme), implemented by lme4 in R (Bates et al, 2014). RT data were regressed onto the interaction between block type (regular block vs. transform block), and cue-target interval length. We included both the linear term and the quadratic term of cue-target interval length since previous studies revealed parabola-shaped RT pattern. Consistent with the random effect structure of previous experiments (Chai et al., 2024, 2025), we also included a random intercept in the model to account for the baseline RT differences between participants (lme formular: RT ∼ (linear cue-target interval + quadratic cue-target interval) * block type + (1 | subject)). The statistical significance of the results was determined and implemented by package lmerTest (Kuznetsova, Brockhoff, & Christensen, 2017). In addition to RT, we also conducted behavioral analyses on accuracy in a logistic regression using the same modelling structure.

### Image acquisition and preprocessing

MRI images were acquired using a 3T scanner (SIEMENS MAGNETOM Prisma), paired with a 64-channel radiofrequency head coil. For the first scanning session, each participant went through an T1-weighted anatomical scan with a MPRAGE sequence (TR = 2250 ms, TE = 4.18 ms, TI = 900 ms, phase encoding direction = AP, acquisition matrix = 256 * 256, FOV = 256 mm, flip angle = 9°, slice thickness = 1 mm, voxel size = 1* 1*1mm). The fMRI images were acquired by using a T2* weighted EPI sequence (TR = 1780 ms, TE = 27 ms, phase encoding direction = AP, image matrix = 84 * 84, FOV = 210 mm, flip angle = 66°, slice thickness = 2.5 mm, voxel size = 2.5*2.5*2.5 mm, multiband factor = 2).

The imaging data were first converted to BIDS format using the BIDScoin toolbox (Zwiers et al., 2022). Subsequently, the data quality was examined by using the MRIQC (23.1.0, Esteban et al., 2017). Several Image quality metrics (IQM) were systematically examined at the group level, including SNR, Framewise Displacement (FD), AFNI’s outlier ratio (aor), AFNI’s quality index(aqi) etc. Following the quality check, the images were preprocessed using the standard fMRIPrep pipelines from fMRIPrep 23.1.0 (Esteban et al., 2019) in a singularity container (Kurtzer et al., 2017). Critically, the functional images went through susceptibility distortion correction (SDC) by using the fieldmap-less approach offered by fMRIPrep to correct for the spatial distortion in EPI images that were caused by magnetic field inhomogeneity. The quality of distortion correction was further visually examined by the main experimenter. All images were spatially normalized to MNI standard space (MNI152NLin2009cAsym) prior to analysis.

### Univariate analysis

#### The main GLM

After preprocessing, the fMRI images were modeled by fitting a whole-brain voxel-wise GLM in SPM12 (www.fil.ion.ucl.ac.uk/spm/software/spm12/) in MATLAB (version R2023a, The MathWorks). For all univariate GLMs, functional images were firstly spatially smoothed (FWHM 8mm) to increase SNR. Since we were interested in task preparation, we mainly focused on the cue-target interval. We included a regressor at the cue-target interval onset for all trials, and another at the start of long cue-target interval window (i.e. 5s after the cue-target interval onset) for all the long trials. In other words, for all short trials, there was one cue-target interval regressor, which was placed at the start of cue-target interval. For all long trials, there were two cue-target interval regressors, one at the start of cue-target interval to capture the activity within short cue-target interval window, and another regressor at 5s after cue-target interval onset to capture the activity within the longer cue-target interval window. The cue-target interval regressors were also further distinguished based on the block type. We additionally included a stimulus onset regressor to account for the variance in BOLD dynamics that is associated with the target image onset. In transform trials, we also included another task cue regressor. We also set separate regressors for trials with incorrect responses. All regressors used stick functions, which were further convolved with the canonical HRF to model the BOLD response. For each run, we further included 10 nuisance regressors, including six motion regressors, a global signal regressor, framewise displacement, CSF signal, and white matter signal.

After fitting the GLM, several contrasts were built for each participant. The main contrast of interest was the difference between regular and transform blocks at the cue-target interval long window. Next, individual contrast images were fed into a second level (i.e. group level) analysis in order to identify brain regions that displayed consistent activation across participants. Significant brain clusters were identified after voxel level (p < .001) thresholding and cluster-level FDR (p < .05) correction.

#### Alternative GLMs

To further evaluated the robustness of our neural findings, we also built three alternative GLMs.

In the first alternative GLM, we excluded all transform trials from the regressors of interest (in contrast to our main GLM where transform instructions were added as separate regressors). This model, naturally, had less statistical power, but helped ensure even more that the difference in visual presentation that followed the cue-target interval in transform trials was not causing any of the observed differences in neural activity between regular and transform trials.

In a second alternative GLM, we used a boxcar function, instead of a stick function, for the cue-target interval regressors in order to model the activation within the whole cue-target interval. All other modeling details were the same as the main GLM. In this alternative GLM, the contrast of interest was the model estimates of BOLD activity within long cue-target intervals between regular and transform blocks. This contrast could not compare the difference only in the second half of the long cue-target interval window because the boxcar function covered the whole cue-target interval, from interval onset to offset. Therefore, this contrast also led to a less sensitive test as we expected the difference between these two block types to arise mainly at the onset of the long cue-target interval window where task uncertainty varied most.

Another GLM was built to further unveil the BOLD dynamics of the significant clusters of brain regions that were identified from the main GLM. The main GLM indicated that several brain regions exhibited stronger activation within the long cue-target interval window in transform versus regular blocks. However, it did not provide information regarding when exactly activation differences occurred and how the BOLD dynamics evolved in each of these brain regions. Because our task is highly dynamic in nature, we aimed to further characterize the BOLD dynamics within these regions. To achieve this, we built another GLM with Finite impulse response (FIR) as the basis function. Specifically, we estimated activation of 10 TRs (1.78s per TR) after task cue onset to obtain one estimate per TR. This time window (i.e. 10 TRs) was chosen after considering both the length of each interval in the task, as well as the lagging nature of BOLD response (Liao et al., 2002). These regressors again distinguished between block types. In order to account for the onset of image stimuli, we included additional stick function regressor at the onset of image stimuli, which were convolved with the canonical HRF. The rest of the GLM was the same as the main GLM.

### Multivariate analysis

#### ROI definitions

Our main regions of interest were the frontoparietal network (FPN) of human brain. The specific ROIs used in the multivariate analyses were selected independently from the univariate analyses, based on a previous study (Assem et al., 2020), which leveraged on a large-scale MRI dataset of 449 healthy subjects from the HCP project. They identified a domain general multiple demand (MD) system by studying contrasts from three cognitively demanding tasks, across 180 areas per hemisphere, as parcellated using the MMP atlas (Glasser et al., 2016). Similar to other previous studies (e.g. Shashidhara et al., 2019; Waskom et al., 2014; Wisniewski et al., 2023), we focused on the FPN, including the anterior prefrontal cortex (aPFC), lateral prefrontal cortex(lPFC), and inferior parietal lobe (iPL) per hemisphere (see the surface plots of Supplementary Figure S4). For full ROI definition, please see Supplementary Table S7.

Similar to previous studies (e.g., Braem et al., 2017; Bhandari et al., 2024), we also chose the auditory cortex as a task neutral network to evaluate the specificity of FPN’s function in representing task information, as the early auditory cortex is not supposed to represent task information. To define this network, we utilized the MMP atlas with its labeling on each parcel, and selected A1, Rl, PBelt, Mbelt, and LBelt per hemisphere as the constituents of the early auditory network.

Finally, to make sure that any results derived from the above ROIs can be generalized to other ROI definitions, we additionally employed an alternative parcellation scheme from Schaefer et al. (2018). This atlas was built by leveraging on both task-fMRI and rs-fMRI across diverse acquisition protocols from 1489 participants, and by adopting a parcellation algorithm that integrate local gradient and global similarity approaches. The granularity of this parcellation ranges between 100 to 1000 parcels, and we used the standard 400-parcel scheme. Each parcel of this atlas is also assigned to a functional network with 17 networks in total that are consistent with Yeo et al. (2011). We selected the FPN ROIs from the Control A and Control B networks. Spatially contiguous parcels were further grouped into ROIs, including aPFC, lPFC, and iPL for each hemisphere, as in the main ROI definition described above. Please see Supplementary Table S8 for the full ROI definition of this alterative ROI definition scheme.

### Task decoding

#### Conjunctive task decoding

a separate GLM was built for the conjunctive task decoding analyses, which was different from the main GLM in two aspects. First, for all multivariate analyses, functional images were not spatially smoothed. Second, all cue-target interval regressors were separately defined based on their conjunctive task identity (one of nine tasks). The resulting t-contrasts of the beta values derived from the first level GLM were used for subsequent decoding analysis.

Task decoding was conducted using the scikit-learn library and packages from Nipype (Gorgolewski et al., 2011) with additional in-house code in Python. Linear support vector machine (SVM) with regularization parameter (*c*) being 1 was employed as the classifier with one-vs-one for multi-class solution. Since we were interested in the difference between block types and between cue-target interval windows, we conducted decoding separately as a function of cue-target interval window and block type. For each condition, we used a leave-one-run-out approach for cross-validation, which means that we trained the classifier using the voxel pattern of a certain brain parcel within a ROI, and their corresponding task labels (1-9) from three runs of data and tested the classifier on the remaining fourth run. We iterated this process by splitting the data into all possible different training and testing sets. The decoding accuracies were then averaged across folds. In the rare case (i.e. 8.14% of total data) of a run with a missing task label (i.e., no examples of that task after data exclusion), the whole run was discarded from cross-validation. The decoding was performed per parcel and per participant.

#### Compositional task decoding

The GLM for compositional decoding used separate task regressors for each task dimension as cue-target interval regressors. Instead of nine tasks as in conjunctive decoding, there were three levels per task dimension (categorization rule versus stimulus type dimension), and thus each cue-target interval window was defined by a combination of two regressors. As in the conjunctive decoding, the resulting t-contrasts of the beta values from the first level GLM were used as input for compositional decoding. Besides these differences, the procedure for compositional decoding was the same as for conjunctive task decoding.

### Representational consistency

We quantified the similarity of voxel patterns of a certain task across runs as a measure of representational consistency. Specifically, for each parcel, the voxel pattern was extracted from the t-contrast maps of the GLM that was built for conjunctive task decoding analyses. Next, for each task separately, we correlated this voxel pattern between every of the six possible between-run combinations (e.g. between run1 and run2, run1 and run3, etc.) using Spearman’s correlation, and the resulting correlation coefficients (i.e. Spearman’s rho) were used as representational similarity index. The higher these coefficients, the more consistent the patterns were. The consistency was computed for each task separately, and then averaged across tasks.

### Statistical inference

The results from the multivariate analysis, including the decoding accuracies and representational similarity indexes, from each participant were fed into a second-level analysis using linear mixed effects models. One statistical model was built for each analysis approach (conjunctive task decoding, compositional, and representational consistency). For task decoding analyses, the lme formula was decoding accuracy – chance level ∼ block type * cue-target interval window * hemisphere * ROI + (1 | subject). That is, the difference between the decoding accuracy and the corresponding chance level (0.1111 for conjunctive decoding, and 0.3333 for all compositional decoding) was regressed onto the interaction between block type (regular versus transform), cue-target interval window (short versus long), hemisphere (right versus left), and ROI (aPFC, lPFC, and iPL). We included hemisphere in the model mainly to capture potential differences in representational strength between ROIs of left and right hemisphere due to our lateralized findings from the univariate results. We also included subject-level random intercepts to account for individual differences in overall decodability, consistent with our current and previous behavioral analyses (Chai et al., 2024, 2025). Another factor of task dimension (stimulus type or task rule) was included as an independent variable in the conjunctive decoding lme. For the representational similarity analysis, the lme formula was representational similarity index ∼ block type * cue-target interval window * hemisphere * ROI + (1 | subject).

In addition to the overall statistical models, we additionally conducted separate analyses for each ROI of interest, namely the aPFC, lPFC, and iPL from the right hemisphere. Since we were mainly interested in examining the interaction between block type and cue-target interval window, a linear mixed effect model was built for each ROI per analysis, with each model including a subject level random intercept. Since one model was built for each pre-defined ROI, no multiple comparison corrections was applied on the statistical results (Rubin, 2024).

## Acknowledgements

This work was supported by an ERC Starting grant awarded to S.B. (European Union’s Horizon 2020 research and innovation program, Grant agreement 852570).

