## Supplementary materials for "Parallel control of conjunctive and compositional representations supports dynamic task preparation"

**Table S1.** The results of linear mixed effect model on participants' task performance (RT)

| Coefficient | Estimate | *SE* | p value |
|---|---|---|---|
| (Intercept) | 1154.5470 | 8.1336 | < .001 |
| Linear CTI | -2408.8130 | 273.0858 | < .001 |
| Quadratic CTI | 762.1240 | 270.1002 | .005 |
| Block type | -20.1409 | 2.4213 | < .001 |
| Linear CTI : block type | -1170.6549 | 273.0581 | < .001 |
| Quadratic CTI : block type | -251.7848 | 270.1286 | .351 |

*Note.* SE = standard error.

**Table S2.** The results of linear mixed effect model on participants' task performance(accuracy)

| Coefficient | Estimate | *SE* | p value |
|---|---|---|---|
| (Intercept) | 2.1997 | 0.0792 | < .001 |
| Linear CTI | 10.8815 | 3.4156 | .001 |
| Quadratic CTI | 3.0257 | 3.3016 | .359 |
| Block type | 0.0408 | 0.0283 | .149 |
| Linear CTI : block type | 1.4734 | 3.3271 | .658 |
| Quadratic CTI : block type | -7.4978 | 3.3639 | .026 |

*Note.* SE = standard error.

**Table S3.** All significant clusters that were identified from the univariate GLM.

| Location | Hemisphere | no. voxels | Peak coordinate | $T_{max}$ | Brodmann |
|---|---|---|---|---|---|
| Transform > Regular | | | | | |
| Lateral PFC (lPFC) | R | 532 | 56, 22.5, 6.5 | 6.2046 | 45 |
| Inferior parietal lobe(iPL) | R | 527 | 53.5, -30, 46.5 | 5.0009 | 40 |
| Anterior PFC (aPFC) | R | 174 | 33.5, 57.5, 1.5 | 5.3682 | 10 |
| Middle temporal gyrus | R | 164 | 48.5, -22.5, -8.5 | 5.3286 | 21 |
| Middle frontal gurus | R | 129 | 33.5, 30, 36.5 | 4.1984 | 9 |
| Regular > Transform | | | | | |
| Parahippocampal Gyrus | L&R | 2845 | -29, -52.5, -6 | -6.6266 | 18 |
| Lateral PFC (lPFC) | L | 494 | -49, 30, 24 | -6.109 | 46 |
| Superior frontal gyrus | L | 201 | -41.5, 10, 56.5 | -6.5068 | 8 |
| Supplementary motor area (SMA) | L | 111 | -4, -10, 59 | -5.0343 | 6 |

**Figure S1.** The BOLD dynamics, which was estimated by using the finite impulse response (FIR) functions in the fMRI GLM, after task cue onset of left Inferior occipital lobe. This ROI was defined anatomically by using the AAL atlas (Rolls et al., 2020).

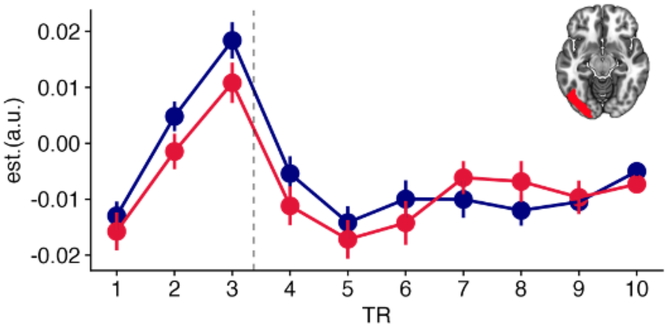

**Figure S2.** Significant voxels (p< .001, w/o cluster level correction) identified by the main GLM (yellow in color) reported in the main text, and an alternative GLM (red in color) which excluded transform trials, with common regions in orange. This alternative GLM was constructed due to a potential confounding factor in the main GLM. In the main GLM, both regular and transform trials from transform blocks were included in the model. However, the trial structure between regular and transform trials were not equivalent since transform trials were featured with another task cue following the CTI (see panel A of Figure 1), which means that image stimuli were shown later in transform trials than in regular trials. Since regular blocks only included regular trials, and transform blocks included both regular and transform trials, any difference found between these two block types can be attributed to the delayed onset of images in transform trials. By comparing the results of these two GLM approaches, the clusters within FPN were consistently identified (in orange color), thus ruling out the potential influence from the confounding factor described above.

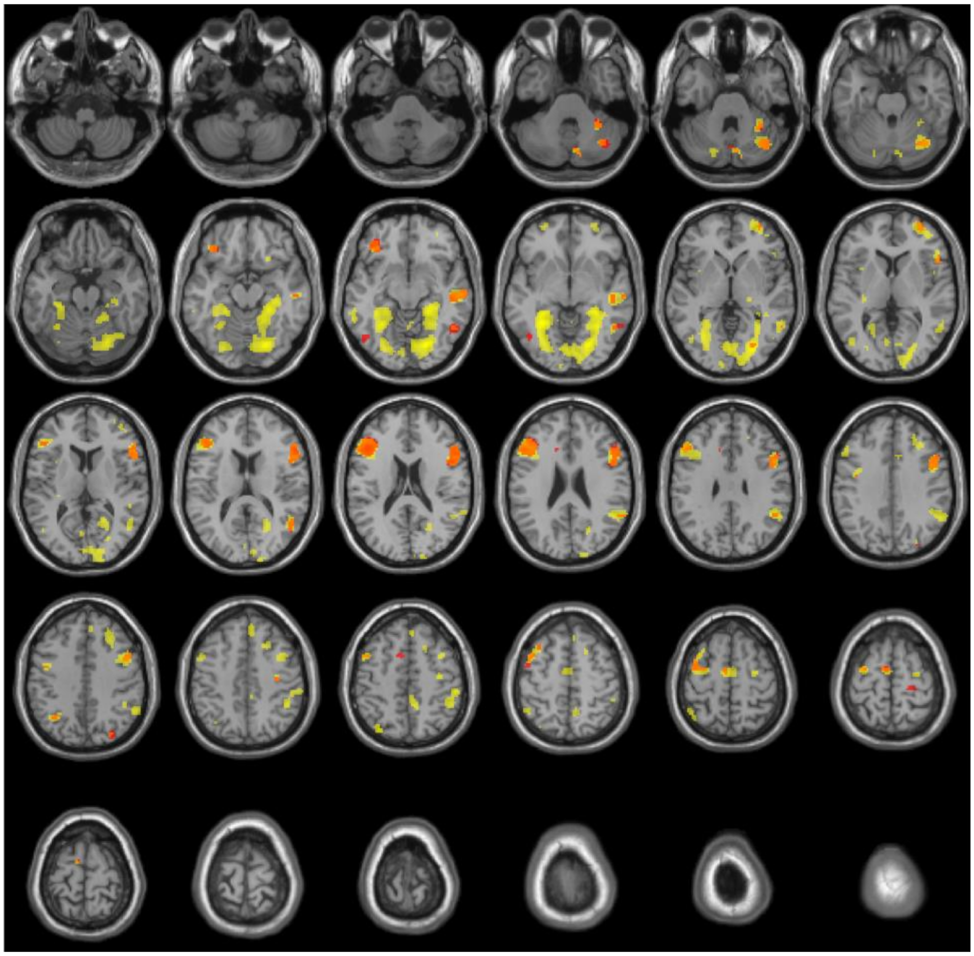

**Figure S3.** Significant voxels (p< .001, w/o cluster level correction), identified by main GLM (yellow in color), and another alternative GLM (red in color), which used boxcar function as CTI regressors, with common results in orange. The contrast of interest from this GLM was the trials of long CTIs in transform blocks versus the ones in regular blocks. the clusters within FPN were also reliably identified by using this alternative GLM approach.

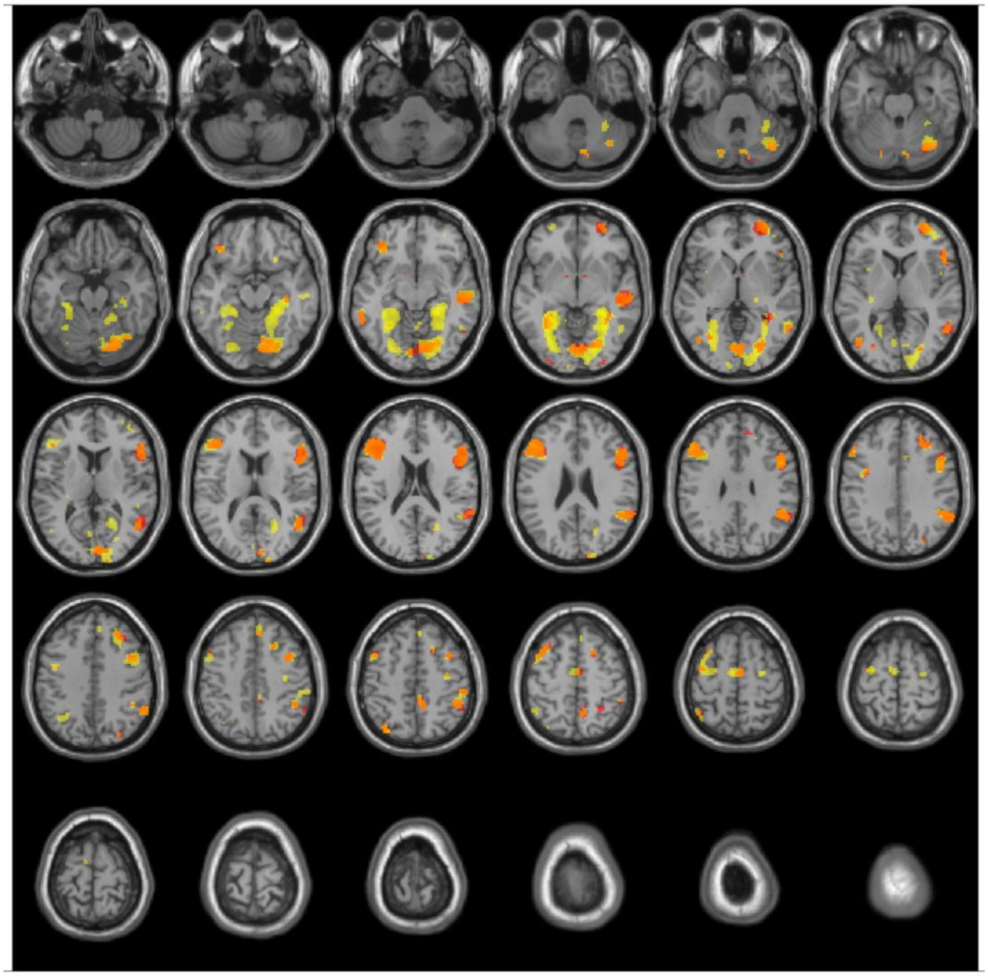

**Figure S4.** The decoding accuracy of conjunctive task representation in the FPN as a function of the interaction between block type and CTI window. dots in the plots denote the mean decoding accuracy across participants, and error bars denote the SE of the mean, grey dashed lines denote the chance-level for conjunctive task decoding (i.e. 0.1111).

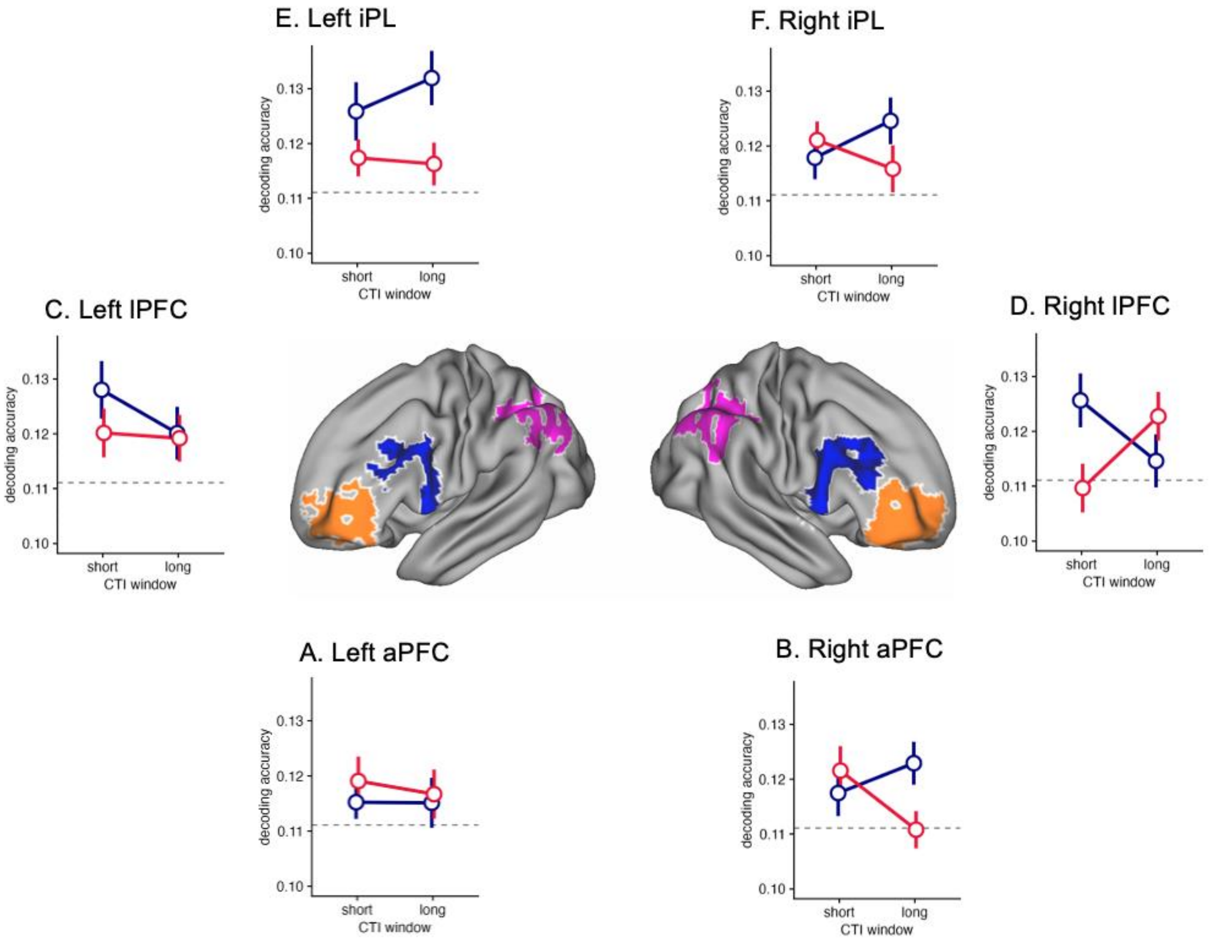

**Table S4.** The results of linear mixed effect model on conjunctive task decoding

| Coefficient | Estimate | *SE* | p value |
|---|---|---|---|
| (Intercept) | 0.0085 | 0.0016 | < .001 |
| block_type | 0.0020 | 0.0007 | .006 |
| CTI_window | 0.0003 | 0.0007 | .647 |
| hemisphere | 0.0008 | 0.0007 | .255 |
| lPFC | -0.0022 | 0.0010 | .031 |
| iPL | 0.0004 | 0.0011 | .700 |
| block_type:CTI_window | -0.0003 | 0.0007 | .713 |
| block_type:hemisphere | 0.0003 | 0.0007 | .735 |
| CTI_window:hemisphere | 0.0002 | 0.0007 | .796 |
| block_type:lPFC | -0.0017 | 0.0010 | .095 |
| block_type:iPL | 0.0000 | 0.0011 | .972 |
| CTI_window:lPFC | 0.0006 | 0.0010 | .538 |
| CTI_window:iPL | 0.0005 | 0.0011 | .653 |
| hemisphere:lPFC | -0.0017 | 0.0010 | .103 |

**Table S4.** The results of linear mixed effect model on conjunctive task decoding

| Coefficient | Estimate | *SE* | p value |
|---|---|---|---|
| hemisphere:iPL | 0.0010 | 0.0011 | .380 |
| block_type:CTI_window:hemisphere | 0.0001 | 0.0007 | .928 |
| block_type:CTI_window:lPFC | -0.0020 | 0.0010 | .047 |
| block_type:CTI_window:iPL | 0.0042 | 0.0011 | < .001 |
| block_type:hemisphere:lPFC | -0.0019 | 0.0010 | .059 |
| block_type:hemisphere:iPL | -0.0001 | 0.0011 | .903 |
| CTI_window:hemisphere:lPFC | -0.0005 | 0.0010 | .594 |
| CTI_window:hemisphere:iPL | 0.0012 | 0.0011 | .303 |
| block_type:CTI_window:hemisphere:lPFC | 0.0017 | 0.0010 | .102 |
| block_type:CTI_window:hemisphere:iPL | -0.0022 | 0.0011 | .052 |

*Note.* The aPFC served as the baseline ROI, SE = standard error.

**Figure S5.** The decoding accuracy of conjunctive task representation in the FPN by using an alternative brain parcellation scheme (Schaefer et al., 2018). ROIs within the FPN were colored in blue. Dots in the plots denote the mean decoding accuracy across participants, and error bars denote the SE of the mean, grey dashed lines denote the chance-level for conjunctive task decoding (i.e. 0.1111).

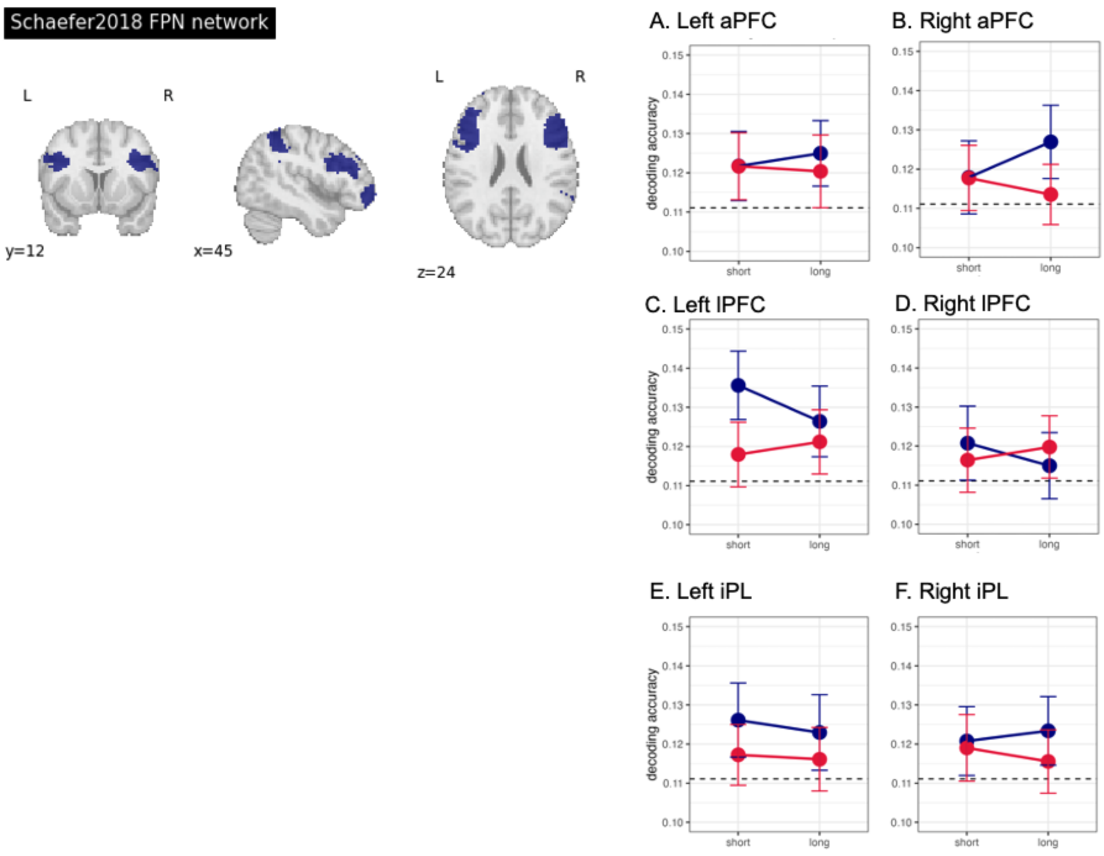

**Figure S6.** The results of compositional task decoding, for both dimension of stimulus type (A-D), and task rule (E-H). A. The mean decoding accuracy of stimulus type in regular and transform blocks. It showed overall above-chance decoding accuracy and higher decodability in regular than in transform blocks. The mean decoding accuracy for each ROI within right FPN as a function of interaction between CTI window and block type was depicted in B. iPL, C. lPFC, and D. aPFC. Across these ROIs, only lPFC exhibit interaction effect which is in line with result from conjunctive decoding. E. The mean decoding accuracy of task rule in regular and transform blocks. Similar to the results of stimulus type decoding, above-chance decoding accuracy and higher decodability in regular than in transform blocks were also observed in task rule decoding. The mean decoding accuracy of task rule for each ROI within right FPN as a function of CTI window and block type was illustrated in F. iPL, G. lPFC, and H. aPFC. No clear interaction effect was observed across ROIs. Noticeably. The compositional decoding accuracy in both task dimensions demonstrated no decreasing trend during task preparation in transform blocks. Bar and dots in the plots denote the mean decoding accuracy across participants, and error bars denote the SE of the mean, grey dashed lines denote the chance-level for compositional task decoding (i.e. 0.3333).

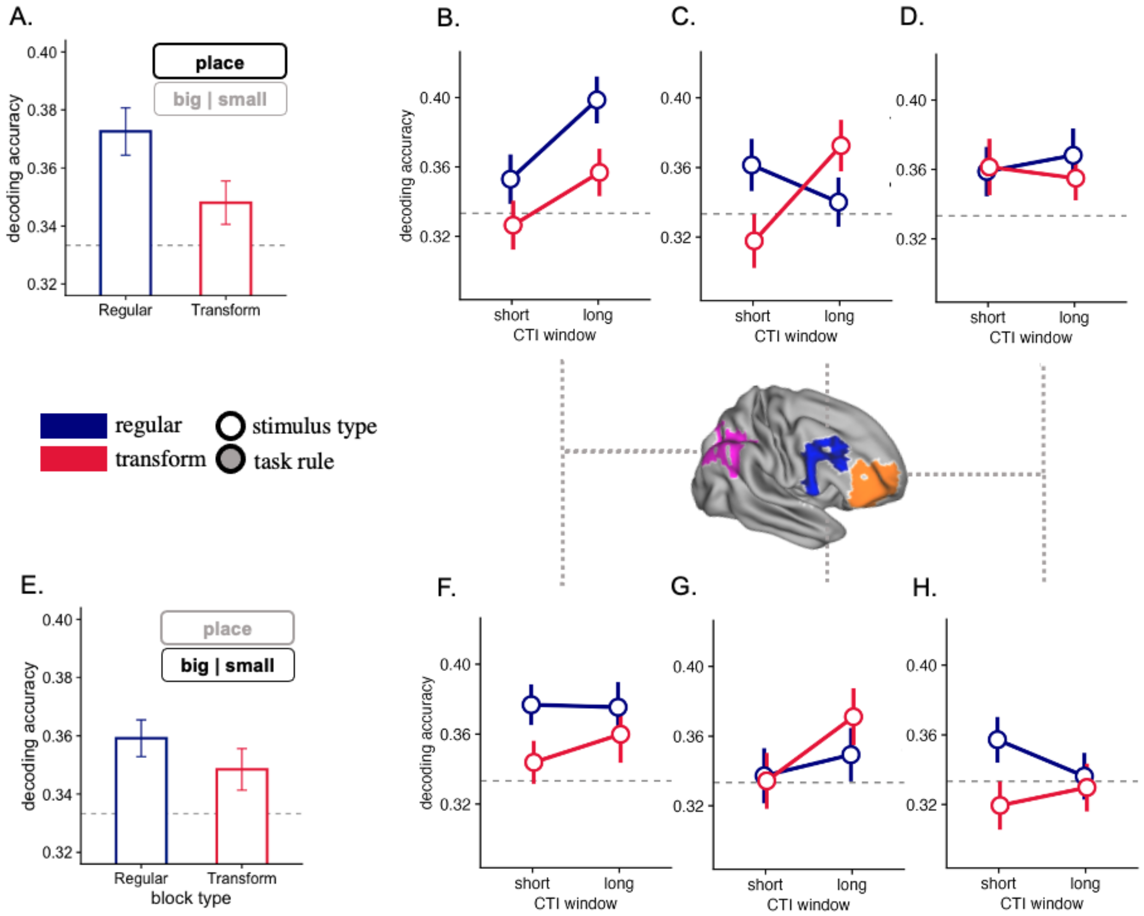

**Table S5**. The results of linear mixed effect model on compositional task decoding

| Coefficient | Estimate | *SE* | p value |
|---|---|---|---|
| (Intercept) | 0.0234 | 0.0037 | < .001 |
| block_type | 0.0082 | 0.0016 | < .001 |
| CTI_window | 0.0046 | 0.0016 | .003 |
| hemisphere | 0.0042 | 0.0016 | .008 |
| lPFC | -0.0079 | 0.0021 | < .001 |
| iPL | 0.0006 | 0.0024 | .814 |
| task_dim | -0.0034 | 0.0016 | .031 |
| block_type:CTI_window | -0.0026 | 0.0016 | .095 |
| block_type:hemisphere | 0.0013 | 0.0016 | .392 |
| CTI_window:hemisphere | -0.0023 | 0.0016 | .137 |
| block_type:lPFC | -0.0045 | 0.0021 | .038 |
| block_type:iPL | -0.0022 | 0.0024 | .349 |
| CTI_window:lPFC | -0.0055 | 0.0021 | .011 |
| CTI_window:iPL | -0.0017 | 0.0024 | .468 |
| hemisphere:lPFC | -0.0036 | 0.0021 | .090 |

**Table S5**. The results of linear mixed effect model on compositional task decoding

| Coefficient | Estimate | *SE* | p value |
|---|---|---|---|
| hemisphere:iPL | 0.0051 | 0.0024 | .032 |
| block_type:task_dim | -0.0036 | 0.0016 | .022 |
| CTI_window:task_dim | -0.0024 | 0.0016 | .116 |
| hemisphere:task_dim | -0.0000 | 0.0016 | .982 |
| lPFC:task_dim | -0.0063 | 0.0021 | .003 |
| iPL:task_dim | 0.0014 | 0.0024 | .551 |
| block_type:CTI_window:hemisphere | 0.0023 | 0.0016 | .133 |
| block_type:CTI_window:lPFC | 0.0001 | 0.0021 | .979 |
| block_type:CTI_window:iPL | -0.0032 | 0.0024 | .177 |
| block_type:hemisphere:lPFC | -0.0045 | 0.0021 | .036 |
| block_type:hemisphere:iPL | 0.0056 | 0.0024 | .019 |
| CTI_window:hemisphere:lPFC | 0.0024 | 0.0021 | .263 |
| CTI_window:hemisphere:iPL | -0.0051 | 0.0024 | .031 |
| block_type:CTI_window:task_dim | -0.0017 | 0.0016 | .288 |
| block_type:hemisphere:task_dim | -0.0029 | 0.0016 | .063 |

**Table S5**. The results of linear mixed effect model on compositional task decoding

| Coefficient | Estimate | *SE* | p value |
|---|---|---|---|
| CTI_window:hemisphere:task_dim | 0.0000 | 0.0016 | .984 |
| block_type:lPFC:task_dim | 0.0025 | 0.0021 | .246 |
| block_type:iPL:task_dim | -0.0015 | 0.0024 | .542 |
| CTI_window:lPFC:task_dim | 0.0015 | 0.0021 | .478 |
| CTI_window:iPL:task_dim | 0.0033 | 0.0024 | .166 |
| hemisphere:lPFC:task_dim | 0.0029 | 0.0021 | .170 |
| hemisphere:iPL:task_dim | -0.0019 | 0.0024 | .424 |
| block_type:CTI_window:hemisphere:lPFC | -0.0029 | 0.0021 | .170 |
| block_type:CTI_window:hemisphere:iPL | 0.0044 | 0.0024 | .062 |
| block_type:CTI_window:hemisphere:task_dim | -0.0005 | 0.0016 | .764 |
| block_type:CTI_window:lPFC:task_dim | -0.0037 | 0.0021 | .086 |
| block_type:CTI_window:iPL:task_dim | 0.0049 | 0.0024 | .041 |
| block_type:hemisphere:lPFC:task_dim | -0.0024 | 0.0021 | .263 |
| block_type:hemisphere:iPL:task_dim | 0.0016 | 0.0024 | .497 |
| CTI_window:hemisphere:lPFC:task_dim | 0.0007 | 0.0021 | .731 |

**Table S5**. The results of linear mixed effect model on compositional task decoding

| Coefficient | Estimate | *SE* | p value |
|---|---|---|---|
| CTI_window:hemisphere:iPL:task_dim | -0.0011 | 0.0024 | .638 |
| block_type:CTI_window:hemisphere:lPFC:task_dim | 0.0010 | 0.0021 | .630 |
| block_type:CTI_window:hemisphere:iPL:task_dim | -0.0027 | 0.0024 | .249 |

*Note.* The aPFC served as the baseline ROI, SE = standard error.

**Figure S7.** The representational consistency, which is quantified by Spearman's rho, in the FPN as a function of the interaction between block type and CTI window. Dots in the plots denote the mean representational consistency across participants, and error bars denotes the SE of the mean.

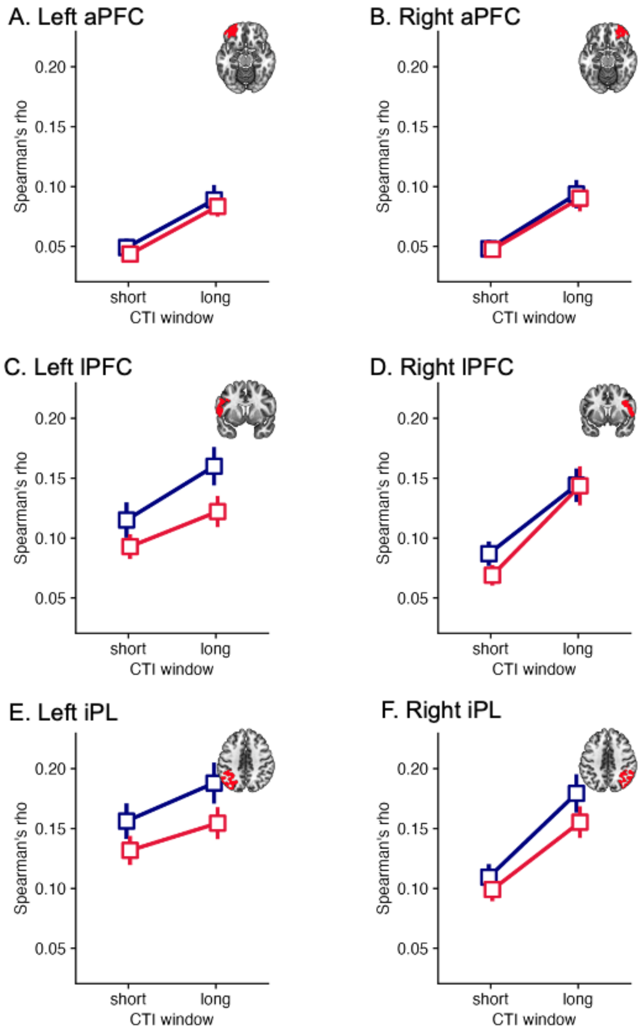

**Table S6.** The results of linear mixed effect model on voxel pattern consistency

| Coefficient | Estimate | *SE* | p value |
|---|---|---|---|
| (Intercept) | 0.1105 | 0.0092 | < .001 |
| lPFC | -0.0424 | 0.0014 | < .001 |
| iPL | 0.0062 | 0.0016 | < .001 |
| hemisphere | 0.0050 | 0.0011 | < .001 |
| block_type | 0.0077 | 0.0011 | < .001 |
| CTI_window | -0.0231 | 0.0011 | < .001 |
| lPFC:hemisphere | -0.0068 | 0.0014 | < .001 |
| iPL:hemisphere | 0.0008 | 0.0016 | .603 |
| lPFC:block_type | -0.0059 | 0.0014 | < .001 |
| iPL:block_type | 0.0021 | 0.0016 | .198 |
| hemisphere:block_type | 0.0030 | 0.0011 | .004 |
| lPFC:CTI_window | 0.0021 | 0.0014 | .142 |
| iPL:CTI_window | -0.0026 | 0.0016 | .100 |
| hemisphere:CTI_window | 0.0058 | 0.0011 | < .001 |

**Table S6.** The results of linear mixed effect model on voxel pattern consistency

| Coefficient | Estimate | *SE* | p value |
|---|---|---|---|
| block_type:CTI_window | -0.0010 | 0.0011 | .348 |
| lPFC:hemisphere:block_type | -0.0021 | 0.0014 | .139 |
| iPL:hemisphere:block_type | 0.0022 | 0.0016 | .171 |
| lPFC:hemisphere:CTI_window | -0.0047 | 0.0014 | .001 |
| iPL:hemisphere:CTI_window | 0.0015 | 0.0016 | .365 |
| lPFC:block_type:CTI_window | 0.0006 | 0.0014 | .660 |
| iPL:block_type:CTI_window | 0.0012 | 0.0016 | .451 |
| hemisphere:block_type:CTI_window | -0.0011 | 0.0011 | .307 |
| lPFC:hemisphere:block_type:CTI_window | 0.0014 | 0.0014 | .327 |
| iPL:hemisphere:block_type:CTI_window | -0.0031 | 0.0016 | .057 |

*Note.* The aPFC served as the baseline ROI, SE = standard error.

**Table S7.** ROIs and their mapping to super parcels from Assem et al.(2020)

| Parcel label | FPN ROI |
|---|---|
| a9-46v | aPFC |
| p47r | aPFC |
| a10p | aPFC |
| 11l | aPFC |
| p10p | aPFC |
| a47r | aPFC |
| 8C | lPFC |
| IFJp | lPFC |
| p9-46v | lPFC |
| 6r | lPFC |
| IP1 | iPL |
| IP2 | iPL |
| PFm | iPL |
| LIPd | iPL |
| MIP | iPL |
| AIP | iPL |
| PGs | iPL |

**Table S8.** ROIs and their mapping to super parcels from Schaefer et al.(2018)

| Parcel label | FPN ROI |
|---|---|
| 17Networks_LH_ContB_PFClv_1 | aPFC |
| 17Networks_LH_ContB_PFClv_2 | aPFC |
| 17Networks_LH_ContB_PFClv_3 | aPFC |
| 17Networks_RH_ContB_PFClv_1 | aPFC |
| 17Networks_RH_ContB_PFClv_2 | aPFC |
| 17Networks_RH_ContB_PFClv_3 | aPFC |
| 17Networks_RH_ContB_PFClv_4 | aPFC |
| 17Networks_LH_ContA_PFClv_1 | lPFC |
| 17Networks_LH_ContA_PFClv_2 | lPFC |
| 17Networks_LH_ContA_PFCl_1 | lPFC |
| 17Networks_LH_ContA_PFCl_2 | lPFC |
| 17Networks_LH_ContA_PFCl_3 | lPFC |
| 17Networks_RH_ContA_PFCl_1 | lPFC |
| 17Networks_RH_ContA_PFCl_2 | lPFC |
| 17Networks_RH_ContA_PFCl_3 | lPFC |
| 17Networks_RH_ContA_PFCl_4 | lPFC |
| 17Networks_RH_ContA_PFCl_5 | lPFC |
| 17Networks_LH_ContA_IPS_1 | iPC |
| 17Networks_LH_ContA_IPS_2 | iPL |
| 17Networks_LH_ContA_IPS_3 | iPL |
| 17Networks_LH_ContA_IPS_4 | iPL |
| 17Networks_LH_ContA_IPS_5 | iPL |
| 17Networks_LH_ContB_IPL_1 | iPL |
| 17Networks_LH_ContB_IPL_2 | iPL |
| 17Networks_LH_ContB_IPL_3 | iPL |
| 17Networks_RH_ContA_IPS_1 | iPL |
| 17Networks_RH_ContA_IPS_2 | iPL |

**Table S8.** ROIs and their mapping to super parcels from Schaefer et al.(2018)

| Parcel label | FPN ROI |
|---|---|
| 17Networks_RH_ContA_IPS_3 | iPL |
| 17Networks_RH_ContA_IPS_4 | iPL |
| 17Networks_RH_ContB_IPL_1 | iPL |
| 17Networks_RH_ContB_IPL_2 | iPL |
| 17Networks_RH_ContB_IPL_3 | iPL |
| 17Networks_RH_ContB_IPL_4 | iPL |
